# Clonal microcolonies recruit individual planktonic colonizers using extracellular matrix factors to assemble *Vibrio cholerae* pellicles

**DOI:** 10.64898/2026.08.20.745993

**Authors:** Hasreet K. Gill, Bonnie L. Bassler

**Affiliations:** Department of Molecular Biology, Princeton University, Princeton, NJ 08544, USA; Howard Hughes Medical Institute, Chevy Chase, MD 20815, USA

**Keywords:** pellicle, gene expression patterning, extracellular matrix

## Abstract

Pellicles, expansive bacterial communities that form on liquid surfaces, contain residents that incorporate asynchronously and exist in distinct physiological states. How interactions between cell populations convert segregated founder pellicle microcolonies into a contiguous, macroscale structure is unknown. Using high resolution timelapse microscopy of *Vibrio cholerae* pellicle formation, we show that the confluent community forms through the recruitment of individual planktonic cells to regions between founder microcolonies. In addition to Type IV MSHA pili, individual planktonic *V. cholerae* cells attach to the air-liquid interface using their cell-surface-bound vibrio polysaccharide (VPS), which binds to the Bap1 and RbmC adhesins secreted by existing pellicle microcolonies. Planktonic *V. cholerae* cells readily attach to the interface by VPS-adhesin binding when they exist in the low-cell-density quorum-sensing state because this is the mode that promotes VPS production. Single-molecule FISH of *V. cholerae* pellicles reveals that regions near pellicle microcolonies, where secreted Bap1 and RbmC adhesin levels are the highest, recruit a higher proportion of VPS-producing planktonic cells than do more distant regions. Thus, existing pellicle microcolony inhabitants in an advanced phase of sessile growth prime the surface for new colonizer cell attachment, driving a spatial pattern of gene expression within the pellicle community.

**SIGNIFICANCE STATEMENT:** Communities of bacteria adhered to liquid-air interfaces exploit phenotypic variation among the constituents to thrive in changing environments. Here, at high spatiotemporal resolution, we show how single cells that join existing communities drive the establishment of particular gene expression patterns across the population. Our findings reveal how bacteria transition from a collection of subcommunities into a contiguous macroscale structure with cells exhibiting a spatially heterogeneous gene-expression pattern.

## INTRODUCTION

Pellicles are microbial communities that reside at air-liquid interfaces and occur widely in medical, environmental, and industrial contexts (1–4). Pellicles form in locations as commonplace as single fluid droplets (5), and as impressive as the ocean surface, where a gelatinous microbial microlayer plays an important role in regulating the global climate (6, 7). In industrial contexts, pellicles have benefited humanity through, for example, their use in vinegar production from antiquity to the present day (8). Pellicles can also be costly and harmful, as in the case of filamentous bulking in wastewater treatment plants (9, 10). Like many naturally occurring microbial communities, pellicles are composed of heterogeneous, multi-species mixtures of bacteria with distinct origins (11–13).

Both Gram-positive and Gram-negative bacterial species form pellicles, and extracellular polymeric substance (EPS) production and flagellar motility are often required for pellicle establishment and maturation (14–23). Imaging studies of pellicle formation, while informative, have largely been carried out at low resolution and/or without longitudinal tracking of live inhabitants (15, 24–26). Thus, the individual cell behaviors that drive pellicle development are not understood. What is known is that incorporation of cells from the planktonic phase occurs during pellicle formation (15, 16, 19, 20, 23, 27, 28). It is not known to what extent planktonic incorporation drives community maturation, and how planktonic cells are recruited to pellicles.

Given the significance of pellicles to humanity and the environment, there is a pressing need for studies to define the mechanisms that drive pellicle maturation. Of particular interest are models to explain the formation of macroscale structures composed of non-clonally-related inhabitants, as this architecture is consistent with microbial communities in the wild. Here, we use the *Vibrio cholerae* pellicle to investigate the microscale events that drive macroscale microbial community formation. Using a method we developed for high resolution confocal imaging at the surface of a thin liquid layer, we demonstrate that the *V. cholerae* pellicle becomes a contiguous structure through incorporation of individual planktonic cells into areas between existing pellicle microcolonies. Focusing on new colonizer recruitment, we demonstrate that in the absence of the Type IV MSHA pili which *V. cholerae* employs for single-cell attachment, binding between vibrio polysaccharide (VPS) on recruited cells and the well-known biofilm adhesins Bap1 and RbmC secreted by pellicle microcolonies enables individual planktonic cells to join the pellicle. Contrary to the established view that cell-surface VPS decoration only occurs in *V. cholerae* cells committed to the microcolony growth mode, we show that planktonic cells are decorated with VPS and this feature is required for Type IV pilus-independent attachment. VPS production is regulated by quorum sensing: the low-cell-density quorum-sensing state promotes, and the high-cell-density quorum-sensing state represses VPS production. Thus, *V. cholerae* planktonic cells in low-cell-density quorum-sensing mode exhibit an attachment advantage over *V. cholerae* planktonic cells in high-cell-density quorum-sensing mode when pili are absent. When *V. cholerae* cells can use both the Type IV pilus-surface interaction and the VPS-adhesin interaction for single-cell attachment, a spatial pattern is generated with higher average *vps* gene expression among cells recruited to regions closest to existing microcolonies. This pattern emerges because VPS-decorated cells joining the pellicle bind to the Bap1 and RbmC adhesins secreted by established microcolonies. The adhesins diffuse away from the producer cells and form a high-to-low concentration gradient with increasing distance from the microcolony edge. Together, our findings reveal how *V. cholerae* redeploys canonical biofilm components during pellicle development to transition from a collection of subcommunities into a contiguous macroscale structure with cells exhibiting a spatially heterogeneous gene-expression pattern.

## RESULTS

### The *V. cholerae* pellicle matures into a contiguous layer via addition of individual planktonic cells in regions between existing microcolonies

To investigate the events that drive pellicle community maturation, we developed a method to culture bacteria in thin liquid layers that permitted single-cell imaging of the air-liquid interface (Figure 1A). We focused on *V. cholerae* because it is a robust pellicle former of global clinical significance, and many reliable tools exist for labeling and genetic manipulation (17, 29, 30). We began by examining the successive steps in wildtype (WT) *V. cholerae* pellicle maturation using a strain carrying chromosomal P*_tac_-ssMBP-AM2.2*, which encodes a constitutively expressed green fluorogen-activating protein (Figure 1B, Movie S1) (31). Real time object tracking during imaging and post-processing to connect objects between frames allowed us to follow individual cells from the time they joined the pellicle plane onward.

**Figure 1.**
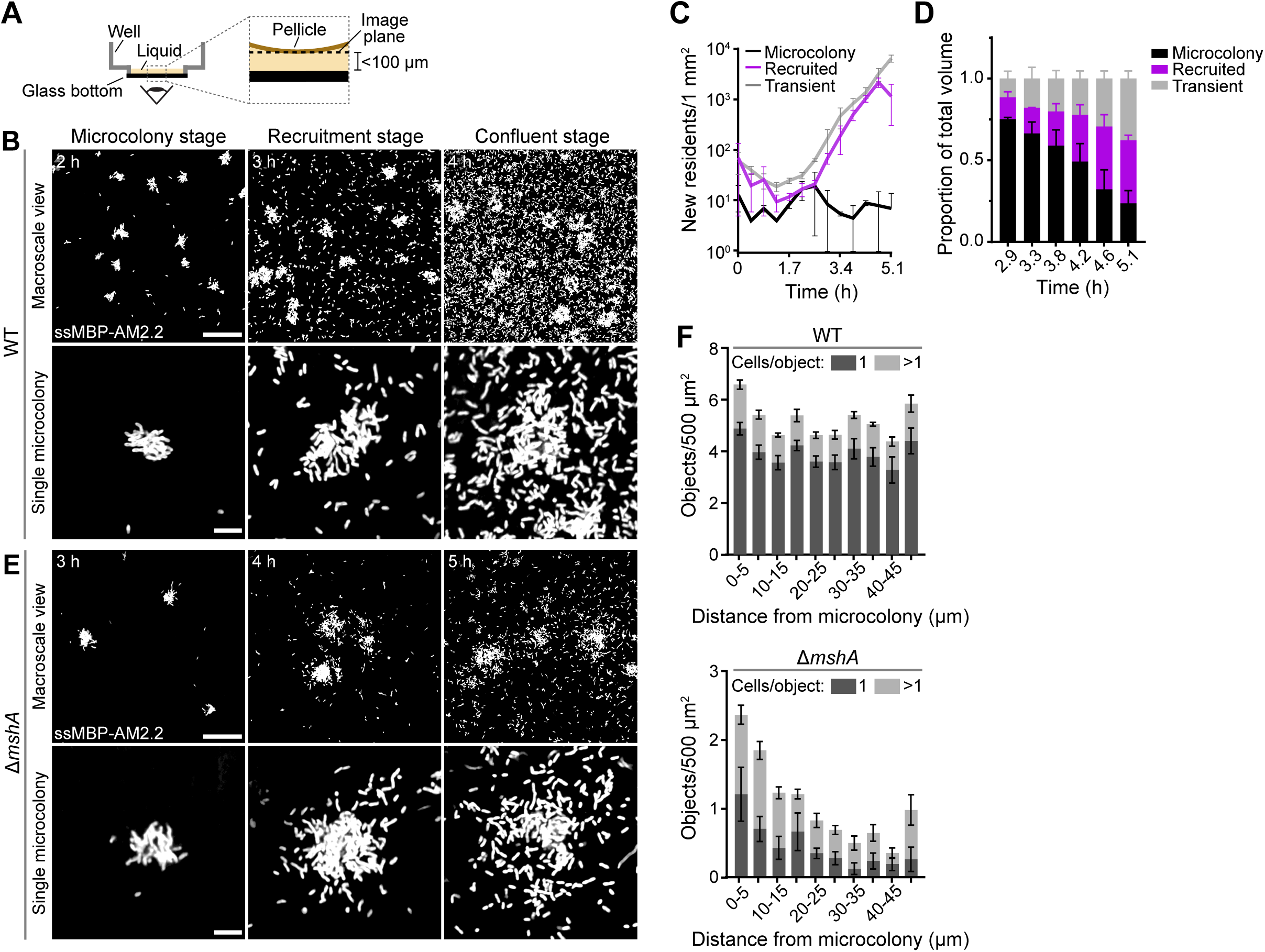
Recruitment of single planktonic cells drives *V. cholerae* pellicle maturation and, in the absence of Type IV MSHA pili, a non-uniform cell density spatial pattern. (A) Schematic representation of the imaging method used to capture timelapse movies and snapshots of *V. cholerae* pellicle development. (B) Snapshots of WT *V. cholerae* pellicle cells constitutively producing ssMBP-AM2.2 taken at single-cell resolution at the air-liquid interface. Time points correspond to distinct stages of pellicle maturation as defined in the main text. (C) Categories of new *V. cholerae* occupants joining the pellicle per time step. (D) Normalized proportion of the total pellicle volume corresponding to each occupant category from panel C during the final 6 time steps. Data in panels C and D are mean values from n=2 replicate timelapse movies. Error bars = SEM. (E) As in panel B for the *V. cholerae* Δ*mshA* mutant. (F) Mean object density separated into single cells (1 cell/object; dark gray) and cell clusters (>1 cell/object; light gray) at the Recruitment stage for WT (3 h, top) and Δ*mshA* (4 h, bottom) *V. cholerae* as a function of distance from the edge of an existing pellicle microcolony. Data are from n=6 replicate single-cell resolution images for each genotype. Error bars = SEM. Ordinary one-way ANOVA (WT: p_1_=0.06, F_1_=1.97, p_>1_=0.19, F_>1_=1.46; Δ*mshA*: p_1_=0.007, F_1_=2.97, p_>1_<0.0001, F_>1_=7.73). Scale bars in panels B and E for the Macroscale and Single microcolony views = 50 μm and 10 μm, respectively.

*V. cholerae* pellicle formation begins with the development of uniformly spaced microcolonies, which attach to the air-liquid interface as single cells and expand clonally while fixed in place (“Microcolony stage”, Figure 1B, Movie S1). Initial microcolonies increase in number from, on average, approximately 10 to 100 within a 1 mm by 1 mm field of view over 5 h (Figure S1A) and, collectively, their volume increases from 60 μm^3^ to 30,000 μm^3^ (Figure S1B). The peak of new cells that arrive to the surface and go on to form microcolonies occurs at about 2 h of culture (Figure 1C, black line), and even at their maximal collective volume at 5 h, initial microcolonies that have undergone three-dimensional growth account for only 20% of the volume of the total resident population at the air-liquid interface (Figure 1D, Figure S1B), showing that the pellicle does not become a contiguous layer through clonal expansion of founder microcolonies.

To understand the composition of the pellicle at maturity before it disassembles due to cell dispersion (“Confluent stage”, Figure 1B), we examined later stages of pellicle maturation and noted the arrival of single cells between established microcolonies. To determine whether these cells originated from microcolonies or from the planktonic phase, we co-cultured otherwise isogenic constitutively red fluorescent (P*_tac_-mScarlet-I*) and green fluorescent (P*_tac_-mNeonGreen*) *V. cholerae* cells and monitored pellicle formation. Red and green cells were highly intermixed in regions between single-color microcolonies, indicating that cells are recruited as non-clonally-related individuals from the liquid phase (Figure S1C, top row). We could define two subpopulations of non-microcolony cells in our timelapse imaging: “Recruited” cells join the pellicle as individuals and persist in place over multiple frames, and “Transient” cells are only present for a single timepoint but are nonetheless temporarily fixed in place and not freely swimming. Tracking of these subpopulations over time revealed that during the growth of the pellicle to confluency, the number of recruited residents and transient single cells on the liquid surface each increase 5-fold with every time step, corresponding to an overall increase of 100-fold each from 3 h to 5 h of culture (Figure 1C, Figure S1A). At the 4 h confluent stage, transient single cells and the descendants of recruited cells account for 50% of the total pellicle volume and increase to ∼75% by 5 h (Figure 1B, D, Figure S1B). A *V. cholerae* pellicle therefore matures into a confluent macroscale community when individual cells from the planktonic phase colonize regions between discrete, clonally-expanded founder microcolonies.

### *V. cholerae* cells lacking Type IV MSHA pili preferentially join pellicles in regions adjacent to existing microcolonies

To understand how interactions between founder microcolonies and subsequent colonizers facilitate formation of the confluent pellicle layer, we focused on the mechanism of single-cell recruitment. *V. cholerae* cells use Type IV MSHA pili to adhere to abiotic solid surfaces (32–37). Extending this understanding to pellicles, we predicted that elimination of these pili might impair attachment of individual recruited colonizers to the air-liquid interface. To test this possibility, we quantified pellicle formation by the *V. cholerae* Δ*mshA* mutant, which does not synthesize Type IV MSHA pili (34, 38). The Δ*mshA* strain carrying P*_tac_-ssMBP-AM2.2* formed microcolonies (Figure 1E) with delayed timing, initiating 1.5 h later than WT *V. cholerae* under our conditions (Figure S1A). The Δ*mshA* mutant formed ∼10-fold fewer founder pellicle microcolonies per 1 mm^2^ than did WT, and the collective volume of the Δ*mshA* mutant was substantially lower than that of WT at the peak of microcolony expansion at (∼4 h, Figure S1B). Although fewer in number, once established, Δ*mshA* microcolonies grew at a rate similar to that of WT microcolonies (Figure S1D). These results are consistent with the loss of Type IV MSHA pili causing diminished, but not abolished, attachment to the air-liquid interface.

The *V. cholerae* Δ*mshA* mutant, like WT *V. cholerae*, showed incorporation of recruited and transient individual cells into the pellicle plane (Figure S1A-B), resulting in a pellicle layer containing single cells and microcolonies at the confluent stage (Figure 1E). As discussed above, individual WT *V. cholerae* planktonic cells that join the maturing pellicle uniformly cover the areas between existing pellicle microcolonies (“Recruitment stage”, Figure 1B). By contrast, individual Δ*mshA* mutant cells concentrate in regions immediately adjacent to established microcolonies (“Recruitment stage”, Figure 1E). Indeed, the local cell density of attached Δ*mshA* single cells decreased over 3-fold from 5 to 30 μm from microcolony edges whereas WT cell density was constant as a function of distance (Figure 1F). Co-culture of red fluorescent and green fluorescent Δ*mshA* cells (Figure S1C, left) showed that, irrespective of distance from the edge of a red or a green microcolony, red and green single cells were equally represented (Figure S1C, right), confirming that, like WT cells, the Δ*mshA* cells are recruited from the planktonic phase and are not microcolony-derived. Together, these results indicate that, during the single-cell recruitment phase of *V. cholerae* pellicle maturation, microcolony cells release a substrate that permits attachment of individual cells via a Type IV MSHA pilus-independent mechanism.

### The *V. cholerae* Δ*mshA* mutant uses binding between VPS and Bap1/RbmC for single-cell attachment to the pellicle

We considered known factors released during biofilm microcolony development as potential candidates for the substrate that recruits individual Δ*mshA V. cholerae* cells to areas around pellicle microcolonies. After a *V. cholerae* cell attaches to a solid surface to initiate biofilm microcolony growth, it produces VPS, which remains attached to the cell that made it (39–41). Biofilm microcolony cells subsequently divide in place and secrete Bap1 and RbmC, which are modular adhesins that bind to and connect VPS on the surfaces of newborn cells to the shared VPS network within the burgeoning microcolony, thereby maintaining clonal growth (42–44). Bap1 and RbmC also adhere the incipient microcolony to the solid surface (45, 46). Thus, the two adhesins can diffuse in-plane away from biofilm microcolony cells that produce them over defined radii (47).

We suspected that individual *V. cholerae* cells lacking Type IV MSHA pili may use binding between Bap1/RbmC secreted by pellicle microcolonies and VPS on their surfaces as a compensatory mechanism to attach and thus enable the mutant pellicle to mature to confluency. To test this possibility, we first added purified Bap1^Δ57aa^ protein to cultures of the *V. cholerae* Δ*mshA* mutant and followed the resulting pattern of cell incorporation into the pellicle. Bap1^Δ57aa^ lacks a hydrophobic loop that tightly associates with abiotic surfaces but retains VPS binding and some surface binding through its β-propeller and β-prism domains, respectively (42, 48, 49). We expected that exogenous Bap1^Δ57aa^ would eliminate the naturally produced gradient of diffused Bap1 adhesin around pellicle microcolonies, effectively rendering all regions of the air-liquid interface equally accessible for *V. cholerae* Δ*mshA* single-cell attachment. Indeed, compared to addition of BSA to the *V. cholerae* Δ*mshA* culture, addition of Bap1^Δ57aa^ increased cell attachment to the air-liquid interface in areas more distal to microcolonies (Figure 2A, Movie S2) and doubled overall pellicle growth measured as the number of static objects in the pellicle over time (Figure 2B). In the *V. cholerae* Δ*mshA* mutant + Bap1^Δ57aa^ condition, the density of single cells residing from 5 to 30 μm away from the microcolony edges decreased ∼1.5-fold compared to over 3-fold in the *V. cholerae* Δ*mshA* mutant + BSA control (Figure 2C). The residual amount of peri-microcolony attachment exhibited by the *V. cholerae* Δ*mshA* mutant in the + Bap1^Δ57aa^ condition likely arises from Bap1 protein that is produced by microcolonies, which we expect adds to the exogenous Bap1^Δ57aa^ protein to increase adhesin concentration close to the microcolony edge. Nonetheless, it is clear that the unvarying presence of a VPS-binding adhesin across the air-liquid interface is sufficient to promote roughly uniform cell recruitment of *V. cholerae* Δ*mshA* mutant cells to the pellicle.

**Figure 2.**
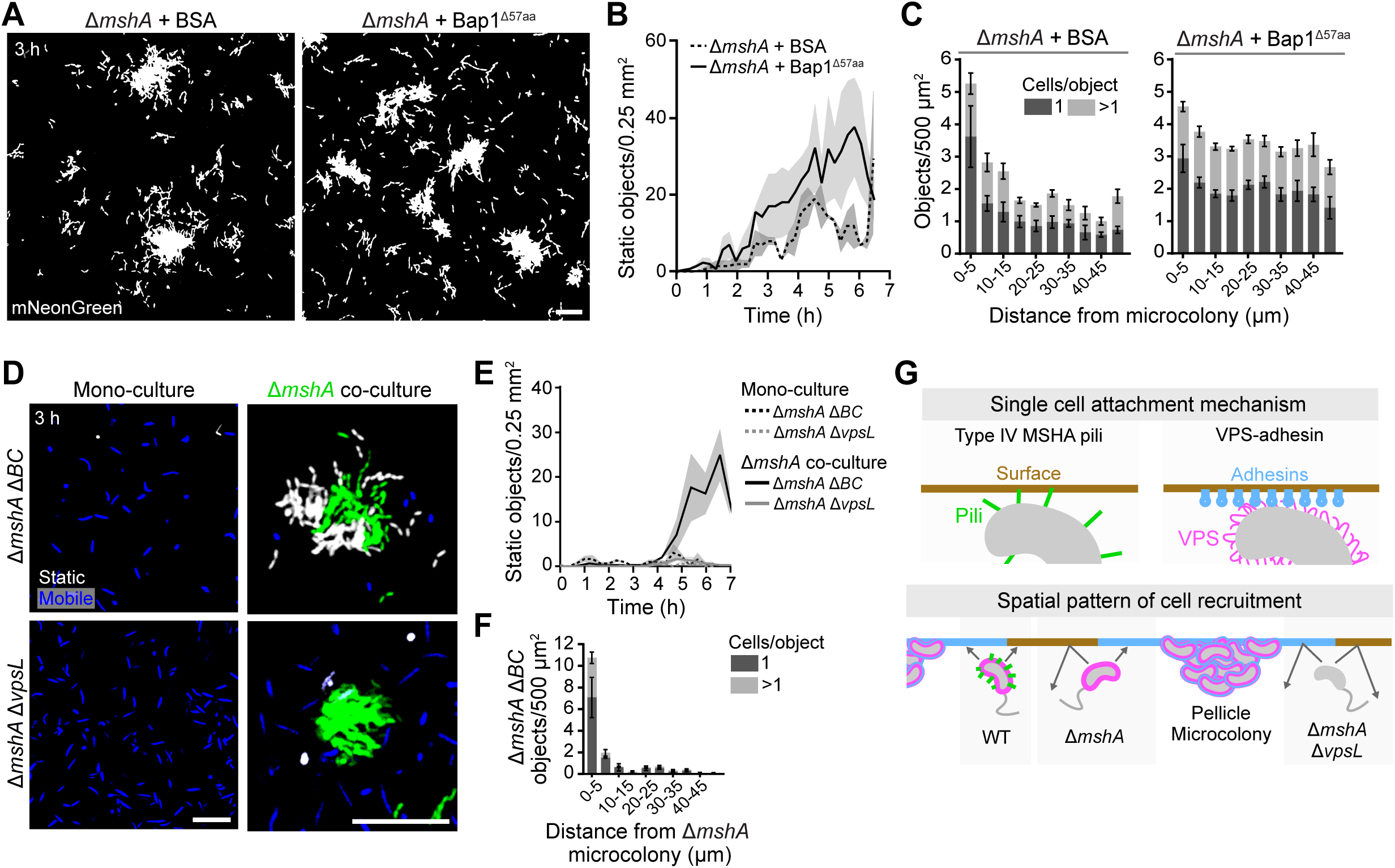
The Bap1/RbmC adhesins secreted by pellicle microcolonies enable recruitment of VPS-decorated Δ*mshA* planktonic *V. cholerae* cells to pellicles. (A) Snapshots as in Figure 1B of Δ*mshA* V*. cholerae* constitutively expressing *mNeonGreen* following addition of 1 mg/mL BSA (left) or 1 μM Bap1^Δ57aa^ (right). (B) Growth of the conditions shown in panel A at the air-liquid interface over time. Data are mean values from n=3 replicate movies taken at high resolution. Shaded error = SEM. Static objects include cells and microcolonies. Comparison of polynomial fits with F-test (p<0.0001; F(DFn, DFd)=16.07(3, 180)). (C) Mean object density as in Figure 1F for the experiments in panel A. Ordinary one-way ANOVA (Δ*mshA* + BSA: p_1_<0.0001, F_1_=6.48, p_>1_=0.0006 F_>1_=3.88; Δ*mshA* + Bap1^Δ57aa^: p_1_=0.01, F_1_=2.63, p_>1_=0.92, F_>1_=0.42). (D) Snapshots of the designated *V. cholerae* mutant strains constitutively expressing *mScarlet-I* in monoculture (left) and co-culture with the Δ*mshA* strain constitutively expressing *mNeonGreen* (right). Mobile and static Δ*mshA* Δ*BC* and Δ*mshA* Δ*vpsL* cells are depicted in blue and white, respectively, and Δ*mshA* cells are shown in green. (E) Growth of the designated *V. cholerae* strains in mono- and co-culture as in panel B for the experiments shown in panel E. Data are from n=2-3 movies per condition/genotype taken at high resolution. (F) As in panels C and D for the indicated *V. cholerae* co-culture condition. (G) Schematic representation of the two modes of *V. cholerae* single-cell attachment to the air-liquid interface during pellicle maturation (top), and the consequence to the spatial patterns of WT and Δ*mshA* single cell incorporation (bottom). In both panels, Type IV MSHA pili are shown in green, Bap1/RbmC adhesions are shown in blue, and VPS is depicted in pink. Tan and blue represent the uncoated and Bap1/RbmC-coated surfaces, respectively. WT *V. cholerae* can attach to both types of surfaces. The Δ*mshA* strain can attach only to the Bap1/RbmC-coated surface near an existing microcolony that secreted the adhesins. The Δ*mshA* Δ*vpsL* strain can attach to neither surface. Scale bars in panels A and D = 20 μm.

To test whether VPS-adhesin binding is required for *V. cholerae* single-cell attachment to pellicles in the absence of Type IV MSHA pili, we generated red fluorescent *V. cholerae* Δ*mshA* Δ*vpsL* mutants, which do not synthesize VPS, and Δ*mshA* Δ*bap1* Δ*rbmC* mutants (we abbreviate Δ*bap1* Δ*rbmC* as Δ*BC*) and followed their pellicle development over time. In contrast to the *V. cholerae* Δ*mshA* mutant, the surface populations of *V. cholerae* Δ*mshA* Δ*vpsL* and Δ*mshA* Δ*BC* mutants were each composed entirely of mobile cells after 3 h of culture (Figure 2D, left) and there was no enrichment of cells at the air-liquid interface relative to the underlying bulk fluid at any point over 6 h (Figure 2E, Figure S2A). To determine whether *V. cholerae* Δ*mshA* Δ*vpsL* and/or Δ*mshA* Δ*BC* cells attached to the interface, but only transiently, we collected short (30 s) high frame rate movies to define the relative proportions of static and mobile cells at the liquid surface. The *V. cholerae* Δ*mshA* mutant had 3% surface-attached cells in accordance with its diminished ability to undergo pellicle microcolony growth, but all *V. cholerae* Δ*mshA* Δ*vpsL* and Δ*mshA* Δ*BC* cells failed to remain at the air-liquid interface for 30 s, resulting in no static cells in either case (Figure S2B). *V. cholerae* cells lacking Type IV MSHA pili therefore require VPS and either Bap1 and/or RbmC to stably bind to and grow at the air-liquid interface.

If *V. cholerae* uses binding between VPS and Bap1/RbmC for single-cell attachment, it follows that *V. cholerae* Δ*mshA* Δ*BC* cells should be capable of binding to exogenously supplied Bap1 and RbmC because they are decorated with VPS, whereas Δ*mshA* Δ*vpsL* cells should not be able to do so. To probe this supposition, we tested whether we could rescue single-cell attachment of the *V. cholerae* Δ*mshA* Δ*BC* and Δ*mshA* Δ*vpsL* mutants by providing an adhesin in the culture. First, we co-cultured red fluorescent Δ*mshA* Δ*BC* or red fluorescent Δ*mshA* Δ*vpsL* cells with green fluorescent Δ*mshA* cells, knowing that the *V. cholerae* Δ*mshA* mutant forms microcolonies (Figure 1E) that secrete Bap1/RbmC. Consistent with the use of VPS-adhesin binding for single-cell attachment, the *V. cholerae* Δ*mshA* Δ*BC* mutant incorporated into the pellicle adjacent to established *V. cholerae* Δ*mshA* microcolonies, including as single cells (Figure 2D, 2F), and proliferated over time within the pellicle (Figure 2E). By contrast, *V. cholerae* Δ*mshA* Δ*vpsL* cells failed to occupy the air-liquid interface (Figure 2D-E). We also supplemented cultures of *V. cholerae* Δ*mshA* Δ*BC* and Δ*mshA* Δ*vpsL* cells with Bap1^Δ57aa^ protein, which rescued pellicle development and uniform cell recruitment of the *V. cholerae* Δ*mshA* Δ*BC* mutant on both short and long timescales but failed to rescue attachment of the *V. cholerae* Δ*mshA* Δ*vpsL* mutant (Figure S2B-D, Movie S3). These results support a model in which, in addition to employing Type IV MSHA pili-mediated attachment, *V. cholerae* single planktonic cells can attach to the air-liquid interface by using their cell-surface VPS to bind to the Bap1 or RbmC adhesin (Figure 2G, top). Since Bap1/RbmC are secreted by microcolonies, VPS-adhesin binding explains peri-microcolony single-cell recruitment during *V. cholerae* Δ*mshA* mutant pellicle formation (Figure 2G, bottom).

### Quorum-sensing status affects Δ*mshA* V. cholerae single-cell attachment near microcolonies

VPS secretion by *V. cholerae* is considered a specific trait of cells residing in biofilm microcolonies, and thus, planktonic cells are not known to be decorated with VPS (50, 51). However, above we have demonstrated that VPS-adhesin binding can be used by planktonic *V. cholerae* single cells for attachment to pellicles. VPS production is regulated by quorum sensing. The prevailing dogma is that, at low cell density, after *V. cholerae* cells have attached to a solid surface using Type IV MSHA pili, the VpsT and VpsR transcription factors activate expression of the *vps* genes (52–56). VPS is made and it promotes biofilm microcolony development. When *V. cholerae* cells on a solid surface reach high cell density, quorum sensing represses *vpsR* and *vpsT*, VPS production is terminated, VPS polymers are removed from cell bodies, and the biofilm disassembles (41, 44, 57, 58). To explain our observations, we hypothesized that, in the planktonic, free-swimming lifestyle, *V. cholerae* cells must produce sufficient VPS when they are in the low-cell-density quorum-sensing state to enable them to incorporate into pellicles by binding to microcolony-secreted Bap1/RbmC. To test this assertion, we measured attachment to the air-liquid interface by planktonic Δ*mshA V. cholerae* cells as function of their culture time and, thus, their population-level of quorum-sensing activation. After growing pellicles of constitutively red fluorescent WT *V. cholerae* to the stage where the air-liquid interface harbors pellicle microcolonies, we spiked-in constitutively green Δ*mshA V. cholerae* cells that had been grown overnight to high cell density, diluted 1:1,000, and pre-cultured from 2 to 7 h (Figure S3A). Our aim was to correlate quorum-sensing state induced by different culture durations with cell attachment ability via VPS-adhesin binding. Our rationale for using the *V. cholerae* Δ*mshA* mutant was to isolate the VPS-adhesin attachment mechanism.

To quantify proficiency of attachment, we calculated the proportion of WT pellicle microcolonies with attached *V. cholerae* Δ*mshA* cells in their peri-microcolony vicinities following spike-in (Figure S3A) and we quantified the data for each pre-culture duration (Figure 3A). We chose the 30 min post-spike-in time point (dashed vertical line in Figure 3A) to discuss here. The attachment trend at 30 min, as a function of pre-culture duration, correlated with expression of the quorum-sensing activated P*_luxC_* promoter fused to *mScarlet-I* (Figure 3B). Regarding the shortest pre-culture duration (2 h), P*_luxC_-mScarlet-I* output was high, consistent with the *V. cholerae* Δ*mshA* cells remaining in the high-cell-density quorum-sensing mode that had been induced during overnight growth. The corresponding fraction of occupied pellicle microcolonies was low, 20% (Figure 3B). With increasing pre-culture duration, P*_luxC_-mScarlet-I* output decreased as *V. cholerae* Δ*mshA* cells transitioned to the low-cell-density quorum-sensing mode (Figure 3B). From 3 h to 5 h of pre-culture, *V. cholerae* Δ*mshA* cell attachment adjacent to microcolonies increased (Figure 3C, Figure S3B, Movie S4), leading to maximum occupancy (80%) of areas around microcolonies by 5 h (Figure 3B). Re-induction of the high-cell-density quorum-sensing state during the 6 and 7 h of pre-culture drove high-level P*_luxC_-mScarlet-I* expression and suppressed the proportion of microcolonies with attached cells to 20% (Figure 3B-C, Movie S4). Half of all attached objects at 30 min post-spike-in following 5 h of pre-culture were single cells and the remainder were clusters of cell doublets or triplets (Figure S3B). These results show that attachment of *V. cholerae* Δ*mshA* single cells adjacent to established pellicle microcolonies correlates with quorum-sensing status: Δ*mshA V. cholerae* cells in low-cell-density quorum-sensing mode show higher attachment than do Δ*mshA V. cholerae* cells in high-cell-density quorum-sensing mode. To directly test the role of quorum sensing in regulation of *V. cholerae* Δ*mshA* single-cell attachment, we generated mutants locked into the high-cell-density and low-cell-density quorum-sensing states in the *V. cholerae* Δ*mshA* parent strain (Δ*mshA luxO^D61A^* and Δ*mshA luxO^D61E^*, respectively). As above, we quantified differences in attachment of these strains after pre-culture for various durations (Figure S3C). Five hours of pre-culture produced maximum attachment of the *V. cholerae* Δ*mshA* and the Δ*mshA luxO^D61E^* strains (Figure 3D, Figure S3C). By contrast, the *V. cholerae* Δ*mshA luxO^D61A^* strain pre-cultured for 5 h showed markedly less peri-microcolony attachment, with only 20% of WT pellicle microcolonies occupied (Figure 3D, Figure S3C). After 7 h pre-culture, the Δ*mshA* and Δ*mshA luxO^D61A^* strains showed minimal peri-microcolony attachment, consistent with both strains being in the high-cell-density quorum-sensing mode, while the locked low-cell-density quorum-sensing Δ*mshA luxO^D61E^* strain showed significantly higher attachment (Figure 3D, Figure S3C). Thus, quorum-sensing status controls the avidity with which Δ*mshA V. cholerae* planktonic cells attach to areas adjacent to pellicle microcolonies.

**Figure 3.**
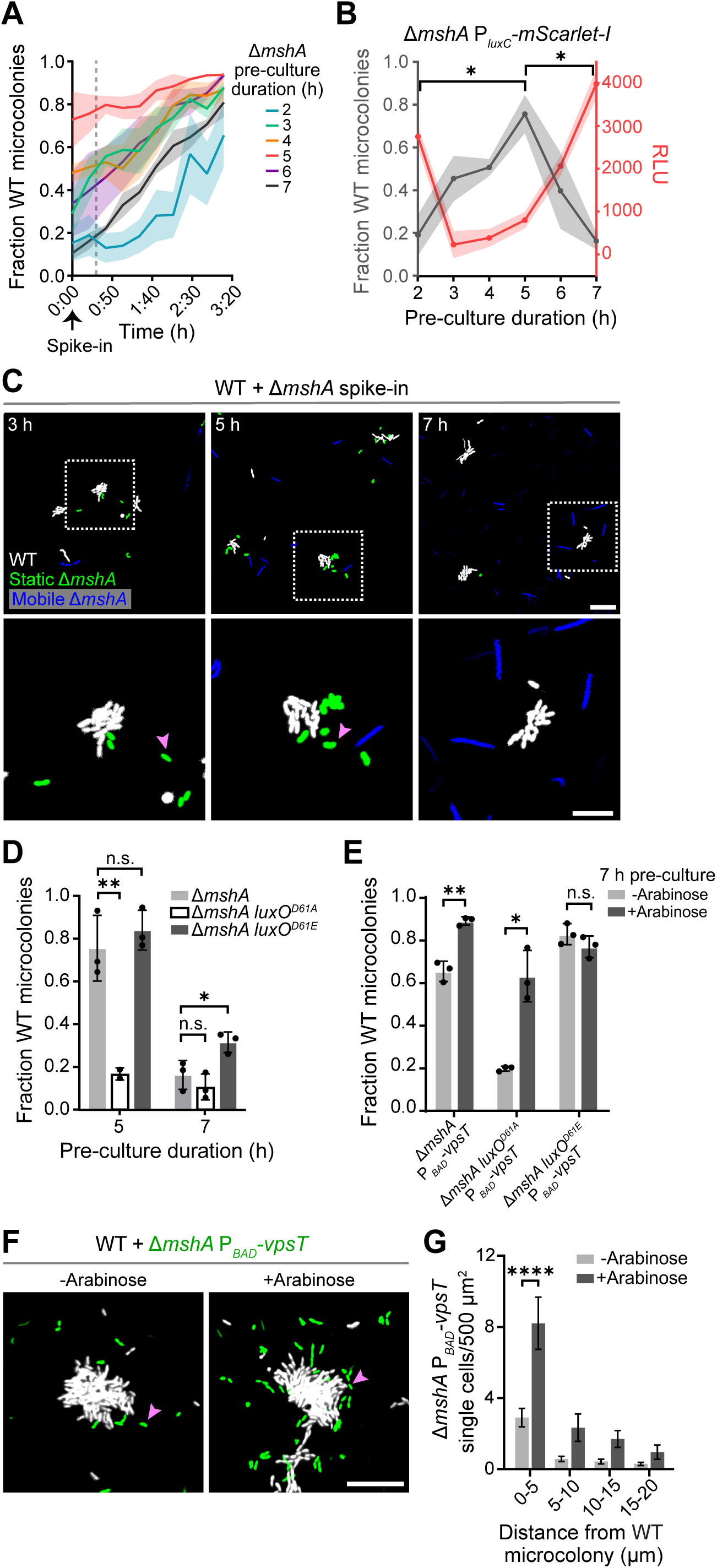
Quorum sensing controls *V. cholerae* planktonic cell attachment to pellicles in the absence of Type IV MSHA pili via regulation of VPS production. (A) Fraction of *V. cholerae* WT microcolonies in which spiked-in *V. cholerae* Δ*mshA* cells occupy their peri-microcolony regions. The Δ*mshA* cells were pre-cultured for the designated times. Curves show the change in mean fraction of occupied WT microcolonies over a short time course imaged at high resolution post-spike-in (arrow). The dashed line indicates the time point used for comparison of pre-culture durations in panel B. Data are from n=3 replicate experiments. Shaded error = SEM. (B) Fraction of occupied WT *V. cholerae* microcolonies from panel A 30 min post-spike-in (gray) following the designated pre-culture durations (*x*-axis). Overlaid is the relative red fluorescence output (red) from the quorum-sensing activated P*_luxC_-mScarlet-I* reporter. Data are from n=3 replicate experiments. Shaded error = SEM. Ordinary one-way ANOVA with Tukey’s MCT performed for Fraction WT microcolonies (p=0.01, F=4.57). (C) Snapshots taken at single-cell resolution 30 min post-spike-in of the *V. cholerae* Δ*mshA* strain into a culture containing WT pellicle microcolonies at the air-liquid interface. Times denote pre-culture durations for the Δ*mshA* strain. Dashed boxes in the top panels are shown enlarged in the bottom panels. Mobile and static *V. cholerae* Δ*mshA* cells are shown in blue and green, respectively. WT cells, static and mobile, are shown in white. (D) Fraction of occupied WT microcolonies as in panel A 30 min post-spike-in for the designated strains pre-cultured for 5 or 7 h. Data are from n=3 replicate experiments. Error bars = SEM. Ordinary one-way ANOVA with Tukey’s MCT (p_5 h_=0.003, F_5 h_=23.21; p_7 h_=0.01, F_7 h_=9.92). (E) Following 7 h pre-culture, fraction of occupied WT microcolonies as in panel A 120 min post-spike-in for the designated strains without (-Arabinose) or with (+Arabinose) induction. Data are from n=3 replicate experiments. Error bars = SEM. Paired two-tailed t-tests. (F) Snapshots as in panel C 120 min post-spike-in of the Δ*mshA* P*_BAD_-vpsT* (green) strain into a culture containing WT pellicle microcolonies (white) at the air-liquid interface without (-Arabinose) or with (+Arabinose) induction. (G) Mean single cell density as in Figure 1F of the *V. cholerae* Δ*mshA* P*_BAD_-vpsT* strain from the experiments in panel F. Two-way ANOVA with Šídák’s MCT (+Arabinose vs. -Arabinose: p<0.0001, F=29.08). Asterisk definitions for Tukey’s/Šídák’s MCT and Paired two-tailed t-test results: ****, p<0.0001; **, p<0.01; *, p<0.05; n.s., not significant. Pink arrows in panels C and F indicate examples of lone Δ*mshA* P*_BAD_-vpsT* cells attached to the air-liquid interface. Scale bars in panels C and F = 20 μm.

### Regulation of VPS production at low- and high-cell density underlies quorum-sensing control of *ΔmshA V. cholerae* single-cell attachment

We predict that *V. cholerae* cells in the low-cell-density quorum-sensing state are primed to attach to the air-liquid interface using VPS-adhesin binding because more cells in the population produce VPS than cells in a population in high-cell-density quorum-sensing mode. We therefore tested whether increasing VPS production in *V. cholerae* Δ*mshA* cells in high-cell-density quorum-sensing mode could restore single-cell attachment to areas adjacent to pellicle microcolonies. To do this, we introduced P*_BAD_-vpsT* into the *V. cholerae* Δ*mshA* strain to conditionally activate *vps* gene expression. We performed our spike-in experiment after 7 h of pre-culture corresponding to *V. cholerae* Δ*mshA* cells exhibiting the high-cell-density quorum-sensing state (Figure 3B). Supplementation with the arabinose inducer drove an increase in Δ*mshA* cell attachment around pellicle microcolonies from 60% to 90% 2 h post-spike-in (Figure 3E-F), with the number of attached single cells increasing at all distances from microcolony edges and roughly tripling in the region nearest the pellicle microcolony, within 5 μm (Figure 3G). We also constructed and performed the same spike-in experiment with the high-cell density locked *V. cholerae* Δ*mshA luxO^D61A^* strain carrying P*_BAD_-vpsT*. A similar increase in cell attachment, from 20% to 60%, occurred (Figure 3E, Figure S3D). The changes in attachment were not a consequence of arabinose supplementation: arabinose addition did not alter cell attachment of the *V. cholerae* strains when they lacked P*_BAD_-vpsT* (Figure S3E). Thus, increasing VPS production fosters single *V. cholerae* cell attachment when cells are in high-cell-density quorum-sensing mode. By contrast, the locked low-cell-density *V. cholerae* Δ*mshA luxO^D61E^* P*_BAD_-vpsT* strain showed no change in attachment with and without arabinose supplementation, indicating that this strain produces sufficient endogenous VpsT to drive maximal VPS-mediated attachment of Δ*mshA* cells to areas adjacent to pellicle microcolonies (Figure 3E, Figure S3D). These results indicate that quorum sensing mediates its effect on the attachment of *V. cholerae* Δ*mshA* cells around pellicle microcolonies by modulating the amount of cell-surface VPS.

### Both Type IV MSHA pili and VPS contribute to the rigidity of single attached *V. cholerae* cells at the air-liquid interface

Our results indicate that *V. cholerae* Δ*mshA* cells can only attach to the air-liquid interface using binding between VPS polymers on their cell bodies and Bap1/RbmC proteins present on the liquid surface. Because WT *V. cholerae* cells have both Type IV MSHA pilus- and VPS-adhesin-driven mechanisms available for single-cell attachment, we wondered if they deploy both mechanisms. We reasoned that *V. cholerae* cells should attach to the air-liquid interface more stably when their surfaces harbor both Type IV MSHA pili and VPS, and when Bap1/RbmC is available compared to when any one of these components is absent. To test this hypothesis, we quantified movement of WT, Δ*mshA*, Δ*BC*, and Δ*vpsL* single *V. cholerae* cells that had already attached to the liquid surface at maximal spatial (<0.1 μm) and temporal (>33 fps) resolution (Figure S4A). Single *V. cholerae* cells lacking Type IV MSHA pili, both Bap1 and RbmC, or VPS exhibit increased mean total distances traveled compared to WT cells (Figure S4B, Movie S5). Moreover, the ratios of the number of cells with long trajectories (unfixed cells) compared to those with short trajectories (fixed cells) are higher for *V. cholerae* Δ*mshA*, Δ*BC*, and Δ*vpsL* single cells than for WT (Figure S4C). These data indicate that, when Type IV MSHA pili are lost or the VPS-adhesin interaction is disrupted, single *V. cholerae* cells are less fixed in place at the air-liquid interface than when both attachment mechanisms are intact, supporting the conclusion that WT *V. cholerae* employs both attachment mechanisms to stabilize new colonization events in the developing pellicle.

### A higher proportion of Δ*mshA V. cholerae* cells recruited to pellicles express *vpsL* than do wildtype recruited cells

Our results support a model in which WT *V. cholerae* can use either or both VPS and Type IV MSHA pili for single-cell attachment to join a developing pellicle, whereas attachment of the Δ*mshA* mutant cell demands decoration with and use of VPS. If so, it follows that the overall proportion of *vpsL-*expressing single cells on the surface should be higher for the *V. cholerae* Δ*mshA* mutant than the WT. To test this supposition, we co-cultured the constitutively green *V. cholerae* Δ*mshA* strain with non-fluorescent WT *V. cholerae* and probed *vpsL* expression among single cells in the resulting pellicles using smFISH (Figure S5A). To capture dynamic changes in gene expression over the course of pellicle maturation, we collected pellicles at both the Recruitment (3 h) and Confluent (5 h) stages (Figure 1B and Figure 4A). *vpsL* expression among the Δ*mshA* single-cell population at the air-liquid interface was significantly different than that for WT single cells at both time points, including after normalization to the housekeeping gene *gyrA* (Figure 4B, Figure S5B). The distributions of per-cell *vpsL* expression were bimodal, with the lower peak corresponding to non-expressing cells. Thus, both the *V. cholerae* WT and Δ*mshA* mutant populations contained *vpsL-*non-expressing and *vpsL-*expressing cells. Our measurements show higher median *vpsL* expression in the Δ*mshA* strain population due to a larger relative number of Δ*mshA* single cells in the maturing pellicle expressing *vpsL* than WT *V. cholerae* cells (Figure S5C). These data track with the requirement for the Δ*mshA* strain to express VPS biosynthetic genes to attach to the liquid surface while WT cells can employ Type IV MSHA pili for attachment, irrespective of *vpsL* expression status. Because *V. cholerae* cells can downregulate *vpsL* expression while retaining cell surface VPS molecules (44), we expect that Δ*mshA vpsL-*non-expressing cells that join the pellicle possess surface VPS despite not actively synthesizing it. Importantly, there was no difference in the relative number of *vpsL*-expressing cells between WT and Δ*mshA V. cholerae* cells in planktonic cultures (Figure S5D, left), suggesting that the difference among pellicle-incorporated single cells stems from higher selective recruitment of VPS-producing Δ*mshA* cells to the air-liquid interface compared to recruitment of WT cells.

**Figure 4.**
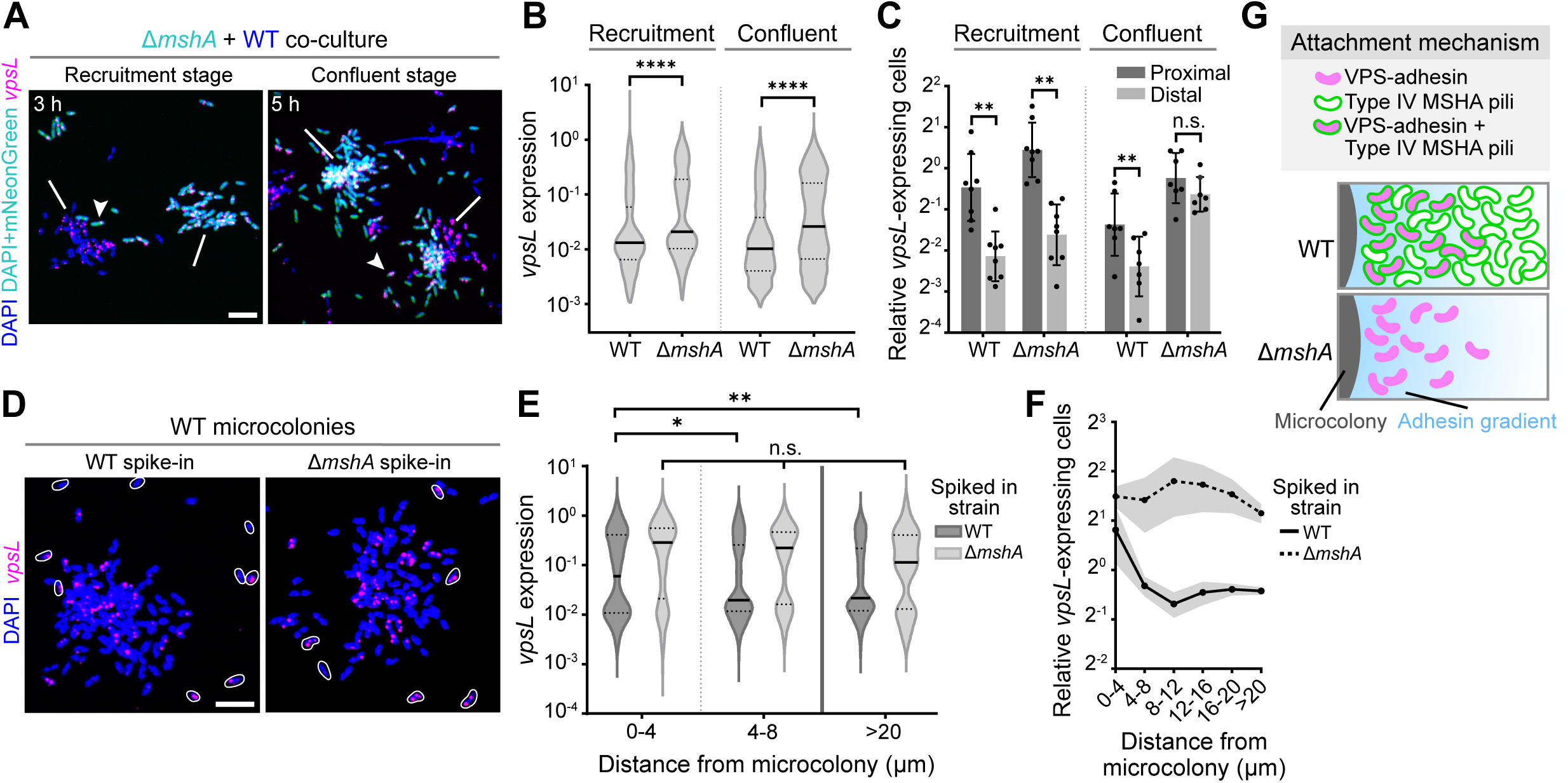
A spatial pattern of *vpsL* expression around pellicle microcolonies forms among recruited *V. cholerae* WT but not *ΔmshA* cells. (A) Maximum projection images of two stages of pellicle growth for co-cultured WT and Δ*mshA V. cholerae*. Images show DAPI (blue), which labels all cells, and smFISH *vpsL* (pink) signals in fixed samples. WT cells, which did not possess a fluorescent reporter, are depicted in blue only, and constitutively *mNeonGreen-*expressing Δ*mshA* cells are depicted in cyan due to the overlay of the blue DAPI and green mNeonGreen signals. Arrows denote examples of single cells expressing *vpsL* and lines denote microcolonies. (B) Distributions of per-cell *vpsL* expression among single cells at the air-liquid interface for the stages shown in panel A. Solid and dotted black lines represent medians and quartiles, respectively. Data are from n=6 replicate co-culture wells per stage (Recruitment: WT, n=1119; Δ*mshA*, n=820; Confluent: WT, n=1665; Δ*mshA*, n=2534). Kolmogorov-Smirnov test (p_Recr._<0.0001, K-S D_Recr._=0.19; p_Conf._<0.0001, K-S D_Conf._=0.21). (C) Proportions of *vpsL*-expressing cells residing proximal (within 20 μm) and distal (further than 20 μm) from microcolony edges for the designated stages in panel A. Data are from replicates shown pooled in panel B (Recruitment: WT_prox._, n=643; WT_dist._, n=476; Δ*mshA*_prox._, n=549; Δ*mshA*_dist._, n=271; Confluent: WT_prox._, n=1128; WT_dist._, n=537; Δ*mshA*_prox_., n=2009; Δ*mshA*_dist._, n=525). Error bars = SEM. Asterisks = paired two-tailed t-tests: ****, p<0.0001; **, p<0.01; *, p<0.05; n.s., not significant. (D) As in panel A for WT microcolonies and the designated spiked-in strains. White outlines show constitutively mNeonGreen-producing spiked-in cells. (E) As in panel B for the cells of the designated spiked-in strains in panel D as a function of distance from microcolonies. Data are from n=6 wells per spiked-in strain (0-4 μm: WT, n=172; Δ*mshA*, n=185; 4-8 μm: WT, n=166; Δ*mshA*, n=157; >20 μm: WT, n=2627; Δ*mshA*, n=51). Kolmogorov-Smirnov tests. (F) Proportion of *vpsL*-expressing cells within 4 μm distance intervals from microcolony edges for the experiment in panel D. Data are from replicates shown pooled in panel E. Shaded error = SEM. (G) Schematic of the region adjacent to a pellicle microcolony illustrating recruited single cell *vpsL* spatial patterning via the mechanism(s) used for attachment, indicated by cell outlines and fill colors. (Top) WT *V. cholerae* uses both Type IV MSHA pili (green outline) and VPS (pink fill) binding to the Bap1/RbmC adhesins (blue gradient), which leads to enrichment of VPS-decorated cells near existing microcolony (dark gray) edges. (Bottom) Δ*mshA V. cholerae* exclusively use VPS binding to the Bap1/RbmC adhesins, leading to high-to-low gradient in cell density driven by the gradient of adhesins secreted from existing microcolonies. Scale bars in panels A and D = 5 μm.

### Recruitment of wildtype *V. cholerae* planktonic cells to pellicles produces a spatial pattern of *vpsL* gene expression around pellicle microcolonies

*V. cholerae* planktonic cells form reversible surface contacts using Type IV MSHA pili that, in time, convert to irreversible binding (33–35). If WT *V. cholerae* can use both the Type IV MSHA pilus- and the VPS-adhesin-mediated mechanisms of single-cell attachment during pellicle maturation, it follows that the subset of VPS-decorated WT cells in the planktonic population may proceed to irreversible attachment more readily in regions coated with Bap1/RbmC than in regions lacking Bap1/RbmC due to VPS-adhesin binding. If so, areas of the developing pellicle surrounding microcolonies, which possess secreted Bap1/RbmC, should preferentially select WT VPS-producing cells from the bulk liquid phase for attachment. To explore this notion, we quantified the spatial distributions of WT *V. cholerae vpsL-*expressing cells that incorporate into pellicles. We defined proximal cells as those located within 20 μm of an existing microcolony and distal cells as those further than 20 μm away. WT *vpsL-*expressing single cells were enriched proximally compared to distally at both the Recruitment (4-fold, Figure 4C) and Confluent (2-fold, Figure 4C) stages, indicating that VPS-producing cells in pellicles indeed become more highly represented in the peri-microcolony regions containing Bap1/RbmC than in more distant regions that are not coated with the adhesins.

To verify that the VPS-producing WT cells responsible for the higher mean *vpsL* expression near microcolonies came from the planktonic culture and not from the microcolonies themselves, we performed smFISH on pellicles where we mimicked the single-cell recruitment phase by spiking green fluorescent WT cells into a non-fluorescent WT culture grown to the microcolony stage (Figure 4D). The planktonic cells that were spiked into the microcolony-containing culture were grown for 5 h beforehand to optimally promote the low-cell-density quorum-sensing state and the highest level of planktonic VPS production (see Figure 3B). Among the spiked-in WT population that attached to the air-liquid interface, the relative population of *vpsL-*expressing cells was larger within 4 μm than in the areas more distant from established Bap1/RbmC-secreting WT pellicle microcolonies (Figure 4E, Figure S5E). The ratio of WT *vpsL*-expressing cells to non-*vpsL*-expressing cells decreased from 2 to ∼0.75 with increasing distance (Figure 4F, Figure S5E). Thus, during WT *V. cholerae* pellicle formation, the Bap1/RbmC gradient surrounding microcolonies can select *vpsL* expressing cells from the planktonic phase, leading to a pattern of high-to-low VPS production among recruited cells proximal-to-distal from existing pellicle microcolonies (Figure 4G).

Regarding attachment of *V. cholerae* Δ*mshA* mutant single cells to pellicles, all attached cells must recently or currently be *vpsL-*expressing. Therefore, we expect no proximal-to-distal *vpsL* spatial pattern as a function of distance from existing pellicle microcolonies. Indeed, while the relative number of Δ*mshA V. cholerae vpsL*-expressing cells versus non-*vpsL*-expressing cells was higher proximally versus distally at the Recruitment stage, this pattern was eliminated by the Confluent stage, consistent with Δ*mshA V. cholerae* cells exclusively using VPS to attach to all locations at the air-liquid interface (Figure 4C). Again, we simulated the Recruitment stage using green fluorescent Δ*mshA* cells spiked into non-fluorescent WT cultures containing pellicle microcolonies (Figure 4D) and, as expected, the spiked-in Δ*mshA* cells showed no difference in mean *vpsL* expression as a function of distance from microcolony edges (Figure 4E). There was a 3- to 4-fold higher proportion of *vpsL*-expressing Δ*mshA* cells than non*-vpsL*-expressing Δ*mshA* cells at all distances (Figure 4F), consistent with overall higher representation of *vpsL-*expressing cells at the surface compared to WT *V. cholerae* (Figure S5D, right). The lack of a *vpsL* spatial expression pattern confirmed the exclusive use of VPS-adhesin binding for air-liquid interface attachment by the Δ*mshA V. cholerae* mutant. Because all instances of cell attachment depend on use of VPS-Bap1/RbmC binding, a pattern of high-to-low cell density is established with distance from the microcolony edge. This pattern reflects the gradient of secreted Bap1/RbmC available for VPS-decorated cells to bind (Figure 1F and Figure 4G).

## DISCUSSION

Large-scale microbial communities have profound consequences for humanity and the planet. These communities often leverage phenotypic variation among resident cells to survive in changing and/or challenging environments (59, 60). To discern how these systems develop spatiotemporal heterogeneity, a mechanistic understanding of their development at single-cell resolution is required (61). Such an understanding would also aid in efforts to predictably construct or modulate the behaviors of such communities. In this regard, the bacterial pellicle represents a tractable model that can be investigated across scales, from molecule to cell to microcolony to macroscale confluent community. *V. cholerae* pellicles harbor cells of unique lineages that are in distinct cell states, with individual cells joining the community at different times during maturation. These features, combined with their thin and sheet-like morphology, which is amenable to high resolution microscopy, make *V. cholerae* pellicles well-suited to the study of higher-order interactions within microbial collectives while still permitting assessment of particular biocomponents at high resolution. In contrast to pellicles, *clonal* biofilm microcolonies have been well studied in *V. cholerae* at high resolution (39, 46, 62–64).

Remarkably, on the scale of a pellicle, the identical components that underpin *V. cholerae* clonal biofilm microcolony development are instead deployed to encourage recruitment of non-clonally related individuals in the regions surrounding pellicle microcolonies. VPS polymers decorate cells within microcolonies as well as a portion of the free-swimming planktonic cells that exist in the low-cell-density quorum-sensing state. The VPS on this subpopulation of planktonic cells fosters air-liquid interface attachment adjacent to microcolonies, where secreted Bap1/RbmC reside. Potentially unrelated colonizers that possess cell surface VPS thus interact with existing residents to drive community growth to confluency. These new colonizers from the planktonic phase are inherently in a different gene-expression regime than cells already present in microcolonies, leading to variations in cell states across the pellicle structure.

It is well established that *V. cholerae* lacking Type IV MSHA pili form biofilm microcolonies with delayed timing (34, 37), and that VPS plays a role (51). However, the attachment mechanism that compensates in the absence of Type IV MSHA pili during biofilm formation has remained undefined. Given our discovery that Δ*mshA V. cholerae* single cells use VPS-adhesin binding for attachment to the air-liquid interface, this same mechanism likely permits solid-surface biofilm attachment of *V. cholerae* Δ*mshA* mutants. If so, the delay in growth known to occur during biofilm microcolony formation (34, 37) and during pellicle formation (this study) is potentially a consequence of free-swimming cells needing to enter the low-cell-density quorum-sensing state so they produce VPS, which permits attachment.

In the case of WT *V. cholerae,* which makes both Type IV MSHA pili and VPS/adhesins, individual cells attach uniformly in the pellicle plane; nonetheless, there is a bias in recruitment of *vpsL*-expressing WT cells to regions nearest existing pellicle microcolonies. We propose that cell surface VPS interactions with Bap1/RbmC-coated regions of the air-liquid interface augment the interactions between Type IV MSHA pili and the air-liquid interface, driving irreversible interface binding (35, 65). Thus, when both attachment mechanisms are operating, VPS-producing WT *V. cholerae* cells gain an attachment advantage. This mechanism is reminiscent of the combined effect of Psl-adhesin and Type IV pili-mediated twitching on single-cell attachment in *Pseudomonas aeruginosa* (66), where initial founders similarly prime the surface for attachment of future colonizers through matrix deposition (67, 68). More detailed studies are needed to measure the individual and combined mechanical contributions of Type IV MSHA pili-surface binding and VPS-adhesin binding during single-cell attachment of *V. cholerae* to pellicles.

Exploitation of diffusible Bap1/RbmC has previously been studied in the context of non-matrix-producing “cheaters” that take advantage of matrix-producing *V. cholerae* biofilm microcolonies (47, 69, 70). Our studies implicate adhesin-sharing as a feature of the normal development of *V. cholerae* pellicles. When viewed on the broad scale of an entire pellicle community, the attachment of VPS-decorated individual cells to areas around established microcolonies may have a beneficial collective outcome. For example, preferential recruitment of *vpsL-*expressing single cells that are in the low-cell-density quorum-sensing state could be a bet-hedging strategy that ensures survival of the broader pellicle community after cells in founder pellicle microcolonies disperse (71). Indeed, extracellular matrix production is costly but advantageous for protection against shear stress and other environmental insults (72, 73). Thus, cells that already produce VPS upon recruitment may promote continued pellicle growth more readily than cells that must initiate VPS production only after reaching the surface. Future studies are required to determine the fates of cells recruited into the pellicle and how their incorporation affects the destiny of the global community.

In the WT *V. cholerae* pellicle, the Bap1/RbmC adhesin gradient establishes a high-to-low spatial pattern of single-cell *vpsL* expression in regions surrounding pellicle microcolonies through selective planktonic cell recruitment. Molecular gradients are used during embryonic development of multicellular organisms to pattern cell states and cell identities. The findings here concerning bacterial pellicle development are reminiscent of eukaryotic development and are therefore consistent with the longstanding view that microbial multicellularity should be considered a developmental process (74–77). Various studies have previously examined gene-expression gradients in macroscale microbial communities; however, they have often focused on fully developed structures, and thus, have rarely captured the spatiotemporal origins of the final gene-expression patterns (59, 78). Here, we reveal the early steps that pattern *vpsL* expression in the developing pellicle at single-cell resolution. While we concentrated on *vps* here, other gene-expression differences could exist between cell types occupying disparate regions of the pellicle, but that possibility has not yet been investigated.

Although the present study concerns maturation of the *V. cholerae* pellicle through recruitment of distinct lineages of the same species, it is possible that the single-cell VPS-adhesin mechanism of attachment also aids in construction of multispecies pellicles in natural contexts. For example, in the sea-surface microlayer (SML), nutrients and polymers adsorbed to the air-ocean interface create a thin, conditioned surface that is used for formation of dynamic microbial communities enriched for *Vibrio* species (6, 79–81). We speculate that multi-species communities like those inhabiting the SML could be established and spatially patterned through VPS-adhesin interactions, e.g., analogous to interspecies aggregation through polysaccharide-adhesin binding during oral biofilm formation (82–84). Indeed, sequence similarities in Bap1/RbmC proteins and VPS biosynthesis enzymes exist broadly across the *Vibrio* genera (85), raising the possibility that cell surface VPS binding to diffusible adhesins could structure the arrangements of single cells and clonal microcolonies from distinct *Vibrio* species in natural pellicles. Mechanistic studies that bridge environmental microbiological approaches with molecular techniques and high-resolution microscopy could probe these possibilities further to discover the breadth and relevance of the findings reported here.

## MATERIALS AND METHODS

### Bacterial strains and molecular methods

All strains used in this study were derived from *V. cholerae* C6706 and are listed in Table S2. Mutants were generated using chitin-induced natural competence and transformation in *V. cholerae* (86, 87). PCR fragments used for transformation that contained mutations of interest were amplified from genomic DNA with at least 2 kb of flanking homology. All mutants were validated by PCR and DNA sequencing.

### Sample preparation for pellicle live imaging

*V. cholerae* pellicles were grown in No. 1.5 glass coverslip-bottomed 24-well plates (MatTek). To establish a thin liquid layer for microscopic imaging, the cell suspension was dispensed directly onto the coverslip surface and spread using a pipette tip to wet the entire glass bottom of the well. For time courses, wells were pre-conditioned with 100 μL of LB medium containing microscopic polystyrene beads used for registration of successive images (TetraSpeck Fluorescent Microspheres, Invitrogen) Beads were subjected to sonication for 10 min followed by a vortex step prior to use. The dish was covered with a gas-permeable seal (Diversified Biotech) and placed at room temperature overnight to permit bead adsorption to the air-liquid interface. For single-cell resolution movies collected using a 60x objective lens, pre-conditioning medium was prepared by diluting 0.2 μm beads 1:50 in LB. For high resolution movies collected using a 20x objective lens, a 1:100 dilution of 0.5 μm beads in LB was used. Wells were washed 5 times to remove excess beads.

For single-cell resolution imaging, overnight cultures were back diluted to an approximate OD_600_ = 0.0002 in LB medium, which were prepared from single colonies inoculated into 5 mL of LB and shaken at 30 °C. For strains expressing *AM2.2*, TO1-2p fluorogen (Bruce Armitage, Carnegie Mellon University) in 100% ethanol/1% acetic acid was diluted 1:1000 into the back diluted cell suspension immediately before plating to achieve a 1 μM final fluorogen concentration. A final volume of 70 μL of cell suspension was added to a pre-conditioned glass-bottom well for confocal imaging. For high resolution imaging, cultures inoculated with single colonies were grown for 4 h with shaking at 30 °C and filtered using 5 μm mesh centrifugal filters (MilliporeSigma) to remove floating cell aggregates. Cultures were back diluted to an OD_600_ = 0.0005 in LB medium and a final volume of 85 μL of cell suspension was added to each pre-conditioned glass-bottom well. For co-culture experiments, equivalent volumes of red and green strains were mixed after back dilution. For experiments involving exogenous addition of Bap1^Δ57aa^ protein or BSA (MilliporeSigma), stocks prepared in 150 mM Tris-HCl were added to achieve the final concentrations indicated in figure legends.

### Confocal imaging of pellicle development

Imaging was performed on a Nikon Eclipse Ti2 inverted microscope equipped with a Yokogawa CSU-W1 SoRa Spinning Disk confocal scanning unit (50 μm pinhole). To mitigate evaporation of thin liquid layers, 20 mL of sterile distilled water was dispensed between wells of the 24-well plate, and the plate was enclosed within a stage-top incubator with a heated lid (28 °C) during imaging (Tokai Hit). Samples were imaged at two resolutions, as indicated in the figure legends. Images taken at single-cell resolution used a CFI Plan Apochromat 60x WI objective lens (Nikon, 1.27 numerical aperture) and those taken at high resolution used a CFI Plan Apochromat Lambda D 20x objective lens (Nikon, 0.75 numerical aperture). A perfluorocarbon and chlorofluorocarbon-based liquid with refractive index 1.33 was used as immersion fluid (Cargille Laboratories). Laser wavelengths of 488 nm (mNeonGreen and AM2.2) and 561 nm (mScarlet-I) were used to excite fluorescent proteins, and the 640 nm wavelength was used to excite TetraSpeck Microspheres. A 0.5 μm *z*-step size was used for all *z*-stacks. Exposure times of 100 ms and 50 ms were used for each channel for 60x and 20x imaging, respectively. Laser power ranged from 8-12% for 60x imaging and 15-20% for 20x imaging. 60x images were taken as tile scans that encompassed 1.0 mm x 1.0 mm regions (5 x 5 tiles), and single-field-of-view images were used for 20x imaging. Samples were imaged for a total of 9 h with approximately 25 min time steps using 60x imaging and 16 h with 20-30 min time steps using 20x imaging.

To image pellicle growth, the air-liquid interface was located and set as the focal plane at the start of each imaging session using adsorbed TetraSpeck Microspheres. Drift in both *xy* and *z* precluded straightforward tracking of imaged cells and microcolonies. We therefore developed a custom macro in the NIS-Elements software that corrected *xy* drift in real time while imaging. Different versions of the macro were developed for 60x imaging and 20x imaging. Briefly, at each time point, the software captured an image of the adsorbed spheres using 640 nm excitation in one field-of-view. For the first time point, the *xy* positions of the spheres were recorded, and for each subsequent time point, the *xy* positions were compared in a pairwise manner to the positions of the spheres recorded at the previous time point. The median *x* and *y* shift values were then applied to the microscope stage controller, which shifted the stage position to correct for drift. Both macros incorporated an Autofocus step at each time point to correct for drift in *z*.

Where indicated in figure legends, snapshots of pellicle development were taken at 60x magnification. In those cases, 100 μL pellicle cultures were permitted to grow as described above until the time point of interest, at which point 30 μL of culture was removed from the well and samples were imaged as *z*-stacks. Unless otherwise indicated, images in figure panels are single *z*-slices.

### High speed imaging of *V. cholerae* cells at the air-liquid interface

To collect high speed and single-cell resolution movies of swimming and attached cells at the air-liquid interface, wells were prepared with 100 uL of overnight cultures back diluted to an approximate OD_600_ = 0.002 after filtration using centrifugal filters as above. Dishes were covered with gas-permeable seals and pellicles were cultured statically at 30 °C until imaging. Single cells at the air-liquid interface were imaged after 1 h of culture, and developing pellicles were imaged after 3 h. Immediately before movie imaging, 30 uL of culture was removed from the well of interest to improve signal quality by decreasing the distance between the sample plane and the objective lens. 30 s movies were captured at the air-liquid interface using a spinning disk confocal microscope (above) equipped with an ORCA-Fusion BT Digital CMOS Camera (Hamamatsu) at 33 frames per s using the “Fast” scanning mode. To capture cell swimming patterns, the 1X magnifier of the SoRa spinning disk unit was used with a CFI Plan Apochromat 60x WI objective lens, while the 2.8X magnifier with a CFI Apochromat TIRF 60x oil objective lens (Nikon, 1.49 numerical aperture) was used to capture gyration of attached single cells.

### Cell spike-in experiment preparation and imaging

Cultures for spike-in experiments were grown overnight as described above. Samples (2 mL) of *V. cholerae* Δ*mshA* strains were prepared every hour by back diluting overnight cultures to OD_600_ = 0.002 in LB. Diluted samples were cultured at 30 °C with shaking until the spike-in experiment was performed. Overnight cultures were maintained under static conditions at room temperature between inoculations. A 2 mL WT culture was grown for 3 h and diluted to OD_600_ = 0.00075 before 100 μL was transferred to each well and grown to the microcolony stage (room temperature for 4 h). OD measurements of pre-cultured samples were used to determine volumes needed to achieve an approximate final OD_600_ of 0.01 in each well for each culture duration. Cultures used for spike-in were diluted directly into wells containing WT pellicles at the microcolony stage and allowed to attach for 10 min before imaging. When needed, arabinose was added at a 0.1% final concentration into both the pre-culture tubes and the imaging wells with the spiked-in cells. An equivalent volume of sterile water was added to controls.

Samples were imaged at the air-liquid interface as described above for 20x high resolution microscopy. However, sphere-based drift correction was not employed, and the built-in ND Acquisition tool was used for time lapse imaging. 10 *z*-slices were taken with a 1.0 μm step size. Each *z*-slice was constructed from a 4 x 4 tile scan to maximize the number of captured pellicle microcolonies. Samples were imaged for 6 h with 20-30 min time steps. Autofocus was performed at the beginning of each time step to correct *z*-drift.

Measurements of P*_luxC_-mScarlet-I* were performed using the same pre-grown samples spiked into WT cultures, prior to dilution. Measurements were taken using a BioTek Synergy Neo2 plate reader. Red fluorescent signal from a blank well containing LB was subtracted from the experimental data, which were subsequently normalized to OD measurements to obtain RFU values.

### Single-molecule RNA-FISH (smFISH) hybridization and imaging

Pellicles were grown for smFISH as described above, but without pre-conditioning with fluorescent spheres. For co-culture experiments with the Δ*mshA* and WT strains, overnight cultures were back diluted to OD_600_ = 0.0002 and OD_600_ = 0.002, respectively. Because WT *V. cholerae* outgrows the Δ*mshA* mutant at the air-liquid interface, 100 μL of the Δ*mshA* strain was plated first and allowed to grow statically at 30 °C for 20 min before 10 μL of WT culture was added, after which the dish was covered with a gas-permeable seal and placed at 30 °C. Recruitment stage pellicles were subsequently grown for an additional 3 h, and Confluent stage pellicles were grown for an additional 5 h. Spike-in experiments used for smFISH were performed as described above.

At the time of sample collection, an 8 mm diameter circular No 1.5 glass coverslip (Electron Microscopy Sciences) was placed on the surface of the liquid to collect the pellicle. After 1 min, 1 mL cold 4% paraformaldehyde in 1X PBS was dispensed under the coverslip for fixation. Samples were fixed for 20 min at room temperature, followed by 4 washes with 1X PBS and 2 washes with LB, after which the floating coverslip was transferred to a new dish containing 70% ethanol in 1X PBS. Samples were permeabilized overnight at 4 °C. Stellaris RNA-FISH probe hybridization was performed as described with minor modifications (50). Volumes were dispensed onto coverslips with adhered pellicles within custom incubation chambers. For mounting, the circular coverslips were placed pellicle side down onto No 1.5 rectangular coverslips with 10 μL Prolong Diamond Antifade mounting medium (Invitrogen) added in between to generate slides for imaging. Slides were cured at room temperature for 2 h and imaged on a spinning disk confocal microscope (above). Planktonic RNA-FISH samples were prepared as described previously (50) and also mounted using 10 μL Prolong Diamond Antifade mounting medium. Sequences of *vpsL-*Quasar 670 and *gyrA-*CAL Fluor Red Stellaris smFISH probe sets can be found in (50).

smFISH samples were imaged using a CFI Apochromat TIRF 60x oil objective lens and the Yokogawa SoRa disk with 2.8X magnification. Samples were excited with the following laser lines: 405 nm (DAPI), 561 nm (CAL Fluor Red), 488 nm (mNeonGreen), and 640 nm (Quasar 670). Images were taken as *z*-stacks with 0.3 μm *z*-step size.

### Statistical methods

Statistical tests were performed using GraphPad Prism. Shapes of underlying distributions were considered when choosing the appropriate statistical test—Kruskal-Wallis tests (used when comparing multiple groups) and Kolmogorov-Smirnov tests (used when comparing the shapes of the cumulative distributions of two groups) were preferred when data were clearly not normally distributed. Otherwise, when comparing replicate means, data were assumed to be distributed normally with similar spreads. Results of statistical tests are reported in figure legends. In cases where asterisks were used to denote p value categories, exact p-values are as follows: Figure S2B (Tukey’s MCT: Δ*mshA* vs. Δ*mshA* Δ*BC,* p=0.04; Δ*mshA* vs. Δ*mshA* Δ*BC* + Bap1^Δ57aa^, p=0.99; Δ*mshA* Δ*BC* vs. Δ*mshA* Δ*BC* + Bap1^Δ57aa^, p=0.01), Figure 3B (Tukey’s MCT: 2 h vs. 5 h, p=0.02; 5 h vs. 7 h, p=0.01); Figure 3D (Tukey’s MCT: 5 h Δ*mshA* vs. 5 h Δ*mshA luxO^D61A^*, p=0.006; 5 h Δ*mshA* vs. 5 h Δ*mshA luxO^D61E^*, p=0.66; 7 h Δ*mshA* vs. 7 h Δ*mshA luxO^D61A^*, p=0.52; 5 h Δ*mshA* vs. 5 h Δ*mshA luxO^D61E^*, p=0.05); Figure 3E (Paired two-tailed t-tests: Δ*mshA* P*_BAD_-vpsT*, p=0.005; Δ*mshA luxO^D61A^* P*_BAD_-vpsT*, p=0.03; Δ*mshA luxO^D61E^* P*_BAD_-vpsT*, p=0.33); Figure 4C (Paired two-tailed t-tests: WT Recruitment, p=0.0047; Δ*mshA* Recruitment, p=0.0035; WT Confluent, p=0.042; Δ*mshA* Confluent, p=0.158); Figure 4E (Kolmogorov-Smirnov tests: WT 0-4 μm vs. 4-8 μm, p=0.05, K-S D=0.15; WT 0-4 μm vs. >20 μm, p=0.003, K-S D=0.14; Δ*mshA* 0-4 μm vs. 4-8 μm, p=0.39, K-S D=0.10; Δ*mshA* 0-4 μm vs. >20 μm, p=0.12, K-S D=0.19). Parameters and outputs for the sum of two Gaussians fits shown in Figures S5C and S5E are reported in Table S1.

## ACKNOWLEDGMENTS AND FUNDING SOURCES

We acknowledge financial support from Howard Hughes Medical Institute (HHMI) and National Science Foundation Award Number MCB-2508324 (B.L.B.). H.K.G. is an HHMI Fellow of the Damon Runyon Cancer Research Foundation (Award Number 2541-25). Purified Bap1^Δ57aa^ protein was generously provided by members of the Rich Olson group, as advised to us by Jing Yan. We are grateful for this contribution. WT *V. cholerae* containing chromosomal P*_tac_-ssMBP-AM2.2* was kindly provided by members of the Andrew Bridges group. We thank Ned Wingreen for his review of the manuscript and members of the Bassler lab for insightful discussions.

## AUTHOR CONTACTS AND COMPETING INTERESTS

Author contacts are as follows: H.K.G. and B.L.B. The authors declare no competing interests.

## AUTHOR CONTRIBUTIONS

H.K.G. and B.L.B. conceived of the study. B.L.B. supervised the study. H.K.G. designed, performed, and analyzed results for all experiments. H.K.G. and B.L.B. wrote the manuscript.

## DATA AND CODE AVAILABILITY

Dataset S1 contains source data underlying all figures. Original scripts are publicly available at https://doi.org/10.5281/zenodo.22016662. Microscopy images and additional information required to analyze data in this manuscript are available upon request from Bonnie L. Bassler.

