## Supplemental Text and Figures for "Clonal microcolonies recruit individual planktonic colonizers using extracellular matrix factors to assemble *Vibrio cholerae* pellicles"

**This PDF file includes:**

Supporting Text

Figures S1 to S5

Tables S1 and S2

Legends for Movies S1 to S5

Legend for Dataset S1

SI References

**Other supporting materials for this manuscript include the following:**

Movies S1 to S5

Dataset S1

### SUPPORTING INFORMATION TEXT

### SUPPLEMENTARY METHODS

#### Data quantitation

Data quantitation was performed using custom MATLAB scripts and ImageJ/FIJI macros. Where indicated in each script for data analysis, code segments were written with assistance from Claude Code (Sonnet 5, Anthropic) provided through Howard Hughes Medical Institute (HHMI).

#### Pellicle timelapse quantitation

For 20x movies, data analysis was performed on the Princeton Della computing cluster. Each *z*-stack per time point was binarized in MATLAB. Binary stacks were analyzed with the TrackMate plugin in ImageJ/FIJI (1), where *z*-position was treated as time, to identify static objects. Plots were generated using all objects or static objects identified by TrackMate as indicated in figures. For the estimated number of cells in Figure S2A, total area per slice was divided by an approximate empirically determined single-cell area.

For 60x movies, several corrections were performed using a custom MATLAB script to track individual cells as they arrived at the air-liquid interface and remained as single cells or grew into pellicle microcolonies. Images were first divided into 4 quadrants to decrease computational load. Images taken of sphere patterns were used to apply minor post-corrections for drift between time points as linear shifts in *x* and *y*. For individual objects, two aberrations were corrected that occurred during *z*-stack acquisition: *xy* drift and stitching artifacts at the boundaries of tiles. For both corrections, a new blank image was created and repopulated with segmented, corrected objects. *xy* drift was corrected by analyzing the positions of fluorescent spheres in each slice and applying *x* and *y* shifts to all objects in the image quadrant per slice. Stitching artifacts were corrected by overlaying all the single-object slices corresponding to the same object in the new, repopulated image. Any objects present in only one *z*-slice, corresponding to unattached or swimming cells, were excluded from image reconstruction. This procedure was performed for each quadrant and for each *z*-stack over time. The TrackMate plugin in ImageJ/FIJI was applied to 2D maximum intensity projections of corrected *z*-stacks over time, and the positions of objects in 3D *z*-stacks were matched to tracked objects in MATLAB.

Plots determining the contributions of pellicle microcolonies and single cells to pellicle growth over time were generated from tracked object data. “Microcolony”, “Recruited” and “Transient” definitions were set as follows: Tracks where the object grew to at least triple its initial volume were designated as Microcolony tracks. If the object’s peak volume never exceeded 150 $\mu m^3$ or if its track lasted for fewer than 5 frames and ended at the final frame of the movie, the track was instead classified as Recruited. Any non-Microcolony track that persisted for at least 3 frames was also classified as Recruited. Any non-Microcolony track that persisted for only 1 frame was classified as Transient. All tracks where the first object’s volume fell below a volume threshold and where the object failed to grow to at least double its initial volume were excluded, as these tracks corresponded to beads used for image registration. Each movie was visually inspected with overlaid track categories to corroborate categorization. We note that the final time step in each movie was excluded from the plots shown in Figure 1C-D and Figure S1A-B because it was not possible to distinguish separate objects. Because of object crowding due to pellicle confluency, at late stages (after 4 h), the small number of new arrivals assigned to the “Microcolony” population are likely objects that merged in subsequent frames.

For snapshots of pellicle development taken at single-cell resolution, the TrackMate plugin in FIJI was used to identify mobile cells in *z*-stacks, where *z* was treated as time. Tracks corresponding to mobile cells were matched to objects in *z*-stacks in MATLAB, and these cells were masked out of full images to generate composite images where mobile cells are represented in the blue channel.

**Spatial pattern quantitation**

Analyses of object positions to assess spatial patterning of pellicle inhabitants was performed using a custom script in MATLAB. *z*-stacks were binarized, and the center *z*-slice for each object was extracted and placed in a new, two-dimensional image used for downstream analysis. Cells only present in a single *z*-slice were excluded. Individual cells were separated from pellicle microcolonies using a size filter. An object was retained as a single cell if the ratio of its long axis to short axis fell within an empirically determined range corresponding to single *V. cholerae* cells (1.2-3), and if it did not divide into two cells upon watershed segmentation. All other objects were considered to be cell clusters. Cells were assigned to distance bins based on the distance from a cell’s centroid to a microcolony edge. The number of cells within each bin were normalized to the total number of pixels found at each distance range.

**High speed movie quantitation**

High speed movies were quantified in ImageJ/FIJI. Background subtraction and binarization were performed, and binarized objects were tracked using the TrackMate plugin. Quantitative properties of resulting trajectories were analyzed in MATLAB. “Mobile” and “Static” categories were defined as follows for Figure S2B: Mobile cells showed a track displacement between 10 and 100 µm and track duration of fewer than 15 seconds, while Static cell tracks were those that lasted for at least 25 seconds with a displacement of less than 10 µm. To identify a cutoff value separating fixed from unfixed cells in Figure S4C, a Gaussian fit was applied to the frequency distribution of WT total distances ($R^2$=0.96, $\mu$=0.9423, $\sigma$=0.50) and the threshold of $\mu+1.5*\sigma$ was chosen.

**Spike-in experiment quantitation**

Analyses of spike-in experiment results were performed in MATLAB and ImageJ/FIJI on the Princeton Della computing cluster. For each *z*-stack, spiked-in cells identified in the green channel were segmented after binarization and tracked in *z* using the TrackMate plugin. Attached cells were identified as those present in the same *xy* position in most *z*-slices. Any attached object that surpassed a size threshold corresponding to a predicted single cell was excluded from further analysis. In MATLAB, pellicle microcolonies in maximum intensity projections of the red channel were segmented after binarization, and, in a manner analogous to the green channel, any object smaller than a size threshold corresponding to a pellicle microcolony was excluded. A pellicle microcolony was considered to possess attached green cells if static cells identified after TrackMate analysis were located within a given area surrounding the microcolony. The area was determined by applying a 100 pixel (approximately 30 μm) dilation of each pellicle microcolony, so all static cells within this region became associated with the pellicle microcolony. Voronoi cell boundaries were generated using the *xy* positions of the centers of pellicle microcolonies and used to separate peri-microcolony areas of adjacent microcolonies that may have merged upon dilation. The fraction of pellicle microcolonies with attached cells was determined after associating static cells in the green channel with pellicle microcolonies in the red channel.

**smFISH quantitation**

smFISH images were analyzed using custom analysis scripts in MATLAB. Maximum intensity projections of *z*-stacks were used for analysis. For co-culture experiments, cells and pellicle microcolonies were segmented using the blue (DAPI) channel. *V. cholerae* Δ*mshA* cells were identified by the mean intensity within each segmented cell in the green channel, which corresponded to signal from constitutively produced mNeonGreen. Microcolonies were identified using a size filter applied to all segmented objects in the DAPI channel. Individual cells were identified from segmented objects in the DAPI channel and retained for further analysis using the method described above for spatial pattern quantitation, which extracted putative single cells on the basis of size and shape. For smFISH analyses of spike-in samples, spiked-in cells were segmented using the green (mNeonGreen) channel, and cells and pellicle microcolonies already present in the culture were segmented using the DAPI channel. This approach permitted identification of spiked-in cells that attached to areas immediately adjacent to pellicle microcolonies, which would have been included as part of the pellicle microcolony if segmentation of green cells had been performed using the DAPI channel. Filtration and exclusion of non-single cells were performed in the same way for co-culture, spike-in, and planktonic smFISH samples.

Quantitation of fluorescent signals corresponding to smFISH target RNAs was performed as described previously, with key changes (2, 3). First, a whole-image background subtraction was performed, and intensity peaks corresponding to smFISH puncta were identified. A 2D Gaussian fit was applied to each peak. To determine gene expression per cell, the sum of the areas under all Gaussian curves within each cell was calculated. For *vpsL* expression, all peak intensities above the background were retained, and cells were determined to be non-*vpsL* expressing if the total fluorescence from that cell fell below a cutoff value determined using a Δ*vpsL* strain control. The Δ*vpsL* sample was grown, prepared, and imaged in an identical manner to WT and Δ*mshA* pellicles. A Gaussian fit was applied to the frequency distribution of log-transformed per-cell signal intensities ($R^2$=0.94, μ=-2.28, σ=0.26) and the threshold value was chosen as μ+3*σ. Thus, negative signals were distinguished on a per-cell basis instead of a per-peak basis. Distances between cells and pellicle microcolonies were determined from the center of a given cell to the edge of the nearest microcolony.

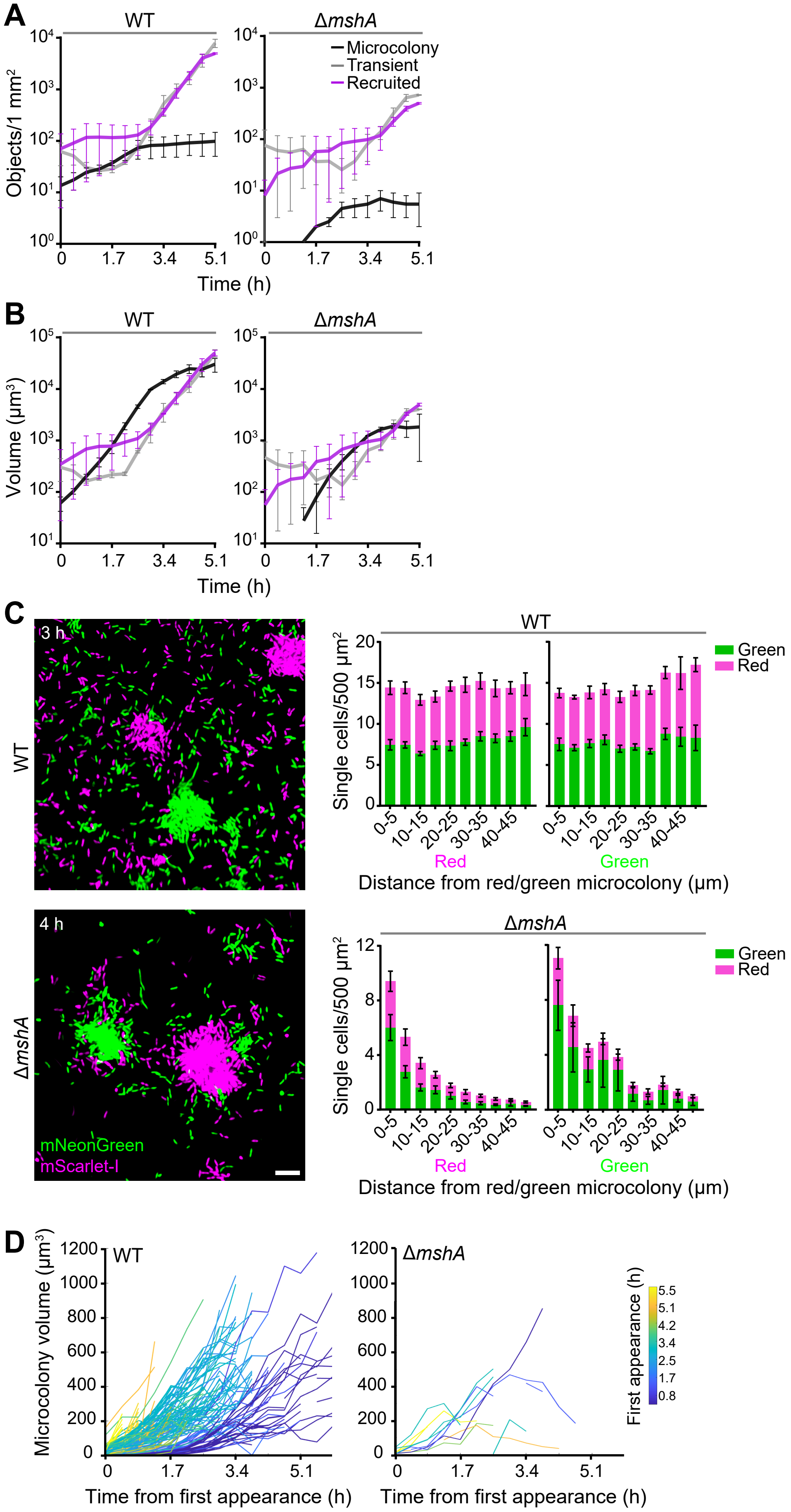


**Figure S1. WT and *ΔmshA V. cholerae* pellicle maturation.**
(A) The total number of objects and (B) total collective volume in *V. cholerae* WT and Δ*mshA* pellicles for the indicated categories of occupants at each time step during pellicle maturation. Data in panels A and B are means calculated from n=2 replicate timelapse movies taken at single-cell resolution. Error bars = SEM. (C) (Left) Snapshots of pellicles following co-culture of the otherwise isogenic WT (top) and Δ*mshA* (bottom) *V. cholerae* strains constitutively expressing *mNeonGreen* (green) or *mScarlet-I* (fuchsia). Scale bar = 10 µm. (Right) Mean density of green and red single cells as a function of distance from a microcolony for the experiments shown in the left panels. Data are from n=5 replicate single-cell resolution images for each genotype. Error bars = SEM. (D) Volumetric growth of individual pellicle microcolonies from timelapse movies taken at single-cell resolution for the WT (n=178 microcolonies) and Δ*mshA* (n=10 microcolonies) strains. Each growth trajectory is aligned to 0 h.

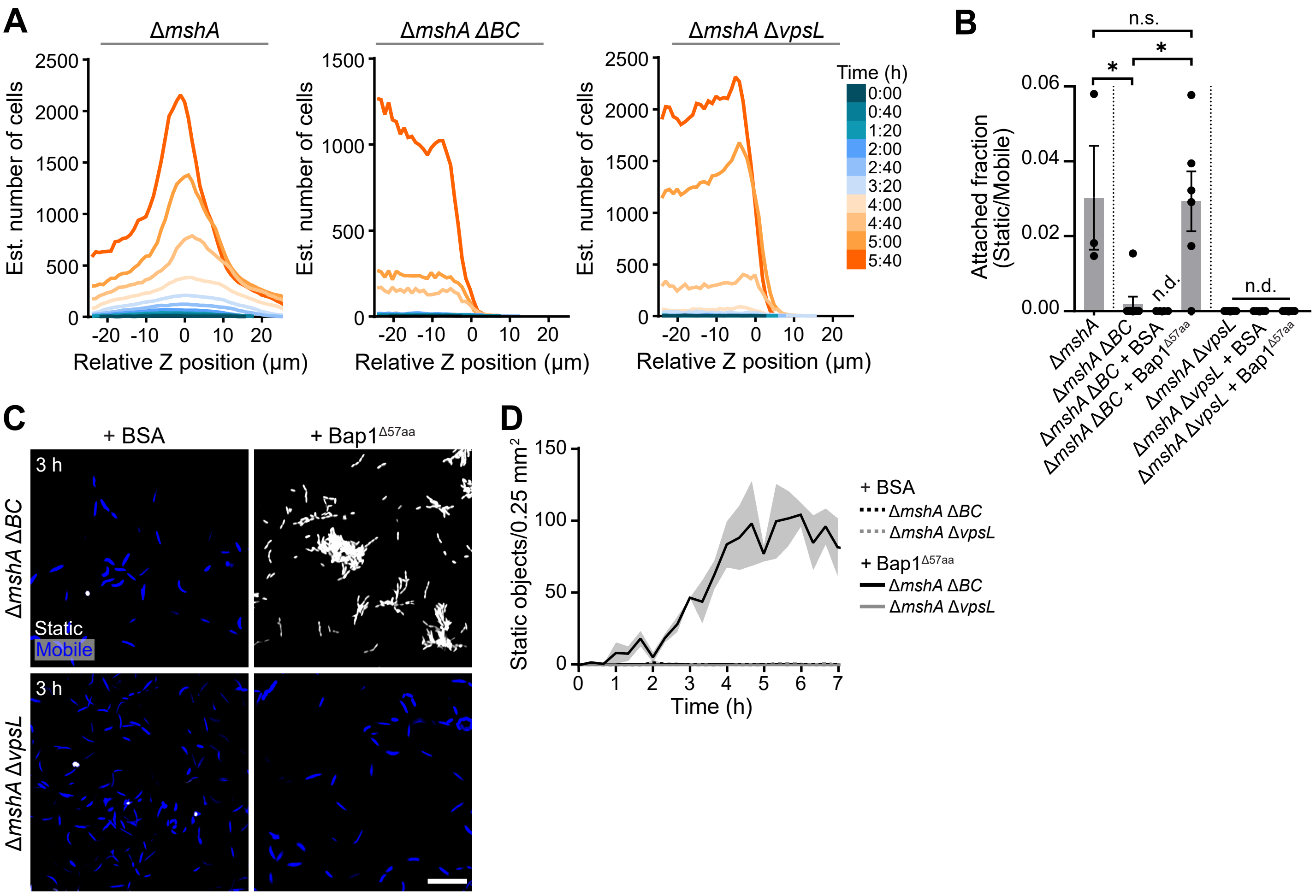


**Figure S2. Growth of *V. cholerae* Δ*mshA*, Δ*mshA* Δ*vpsL*, and Δ*mshA* Δ*BC* mutants at the air-liquid interface.**
(A) Growth of representative cultures of the designated strains imaged at high resolution over time as a function of z-position, where 0 µm corresponds to the air-liquid interface. (B) Fraction of attached cells of the designated strains determined from 30 s high frame rate movies taken at the air-liquid interface. Data are from n=3-6 replicate movies. Error bars = SEM. Ordinary one-way ANOVA with Tukey's MCT ($p=0.008$, $F=6.95$; *, $p<0.05$; n.s., not significant; n.d., not detected). (C) Snapshots at the air-liquid interface for the designated strains that constitutively produce mScarlet-I with the designated proteins added. Mobile and static cells are indicated in blue and white, respectively. Scale bar = 20 µm. (D) Growth of the designated strains at the air-liquid interface over time corresponding to conditions in panel C. Static objects include individual cells and microcolonies. Data are from n=2 timelapse movies per condition/genotype taken at high resolution. Shaded error = SEM.

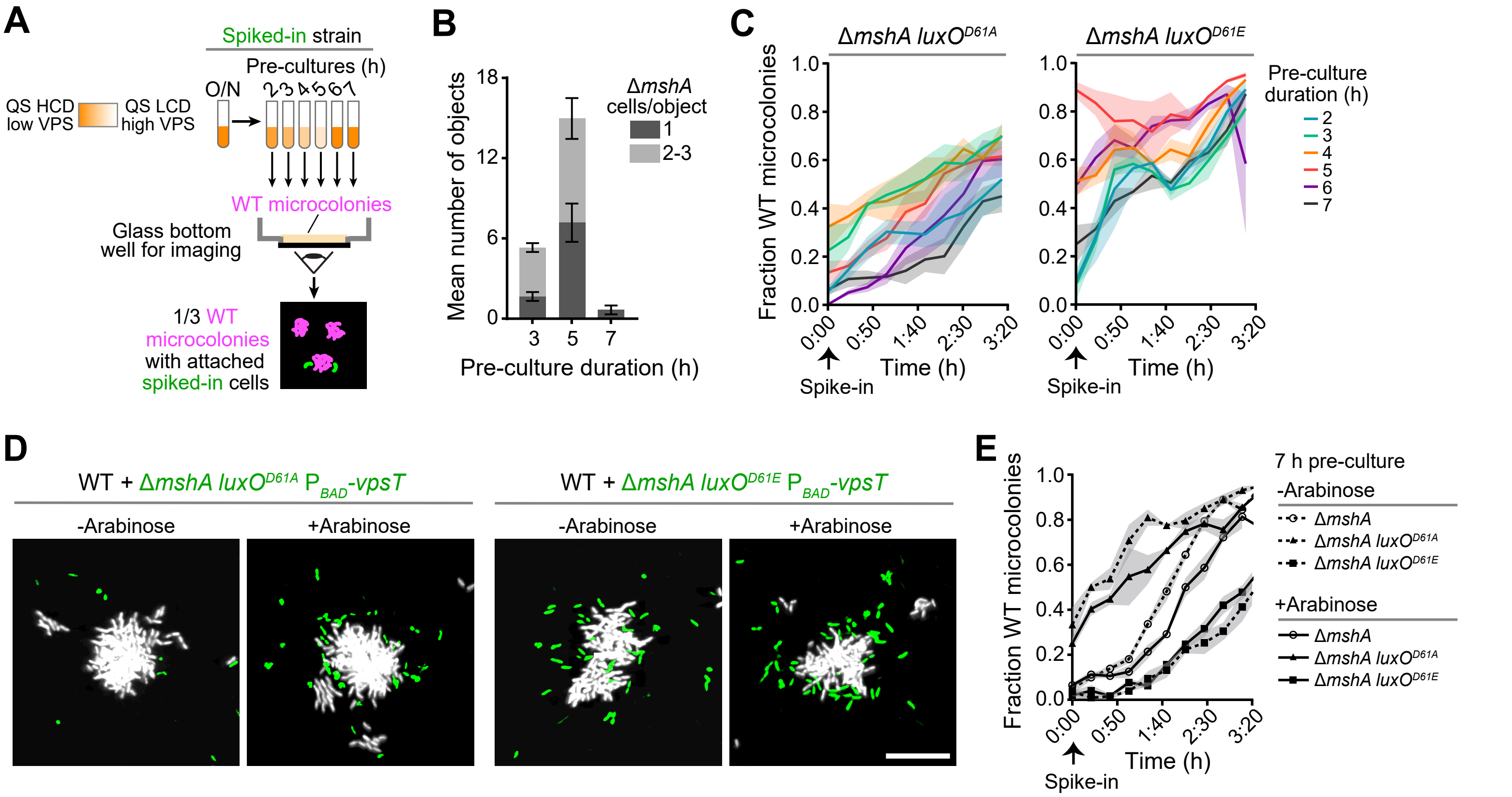


**Figure S3. *V. cholerae* Δ*mshA* single cell recruitment is controlled by quorum-sensing regulation of VPS production.** (A) Schematic representation of the experimental procedure for spike-in experiments used throughout this work. QS, HCD, LCD, O/N designate quorum sensing, high-cell density, low-cell density, and overnight, respectively. (B) Mean number of objects attached to the air-liquid interface 30 min post-spike-in for the Δ*mshA* strain and designated pre-culture durations. Data are from n=3-5 replicate single-cell resolution images per pre-culture duration. Error bars = SEM. (C) Fraction of *V. cholerae* WT microcolonies with the designated spiked-in cells occupying their peri-microcolony regions as in main Figure 3A. Data are from n=3 replicate experiments. Shaded error = SEM. (D) Snapshots as in main Figure 3F 120 min post-spike-in of the designated $P_{BAD}$-*vpsT*-expressing strains (green) into a culture containing WT pellicle microcolonies (white) at the air-liquid interface without and with Arabinose induction. Scale bar = 20 μm. (E) As in panel C after 7 h pre-culture for the designated strains without and with Arabinose induction. Data are from n=3 replicate experiments. Shaded error = SEM.

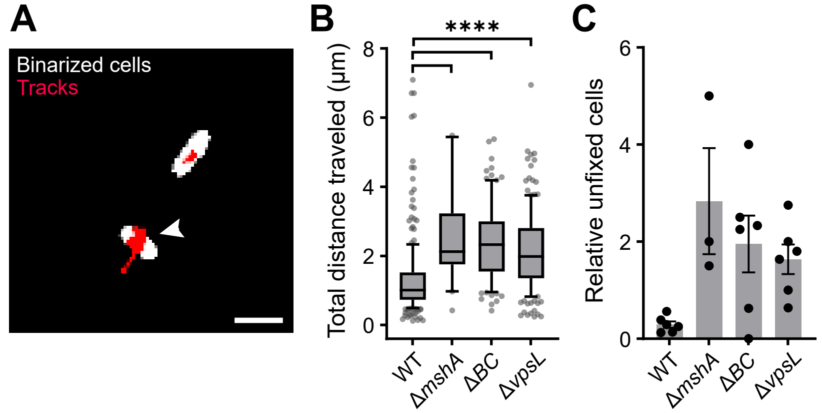


**Figure S4. Rigidity of WT, Δ*mshA*, Δ*BC*, and Δ*vpsL V. cholerae* single-cell attachments to the air-liquid interface.** (A) Binarized frame from a 30 s high frame rate and single-cell resolution movie of attached cells at the air-liquid interface for the *V. cholerae* Δ*vpsL* mutant constitutively expressing *mScarlet-I*. Tracks (red) connect positions of cell centroids across all frames, thus representing movement as spread in X and Y axes. Arrow denotes an unfixed cell. Scale bar = 1µm. (B) Total distances traveled as judged by single-cell tracks for the designated strains. Data are from n=3-6 replicate movies per genotype (WT, n=236; Δ*mshA*, n=20; Δ*BC*, n=98; Δ*vpsL*, n=160). Whiskers, 10-90th percentile. Kruskal-Wallis test (p<0.0001, K-W statistic=115.3; ****, p<0.0001). (C) Ratio of total distances traveled above vs. below an empirically-determined cutoff value (unfixed to fixed cells) for the designated strains calculated per replicate for the data in panel B. Error bars = SEM. Ordinary one-way ANOVA (p=0.02, F=4.28).

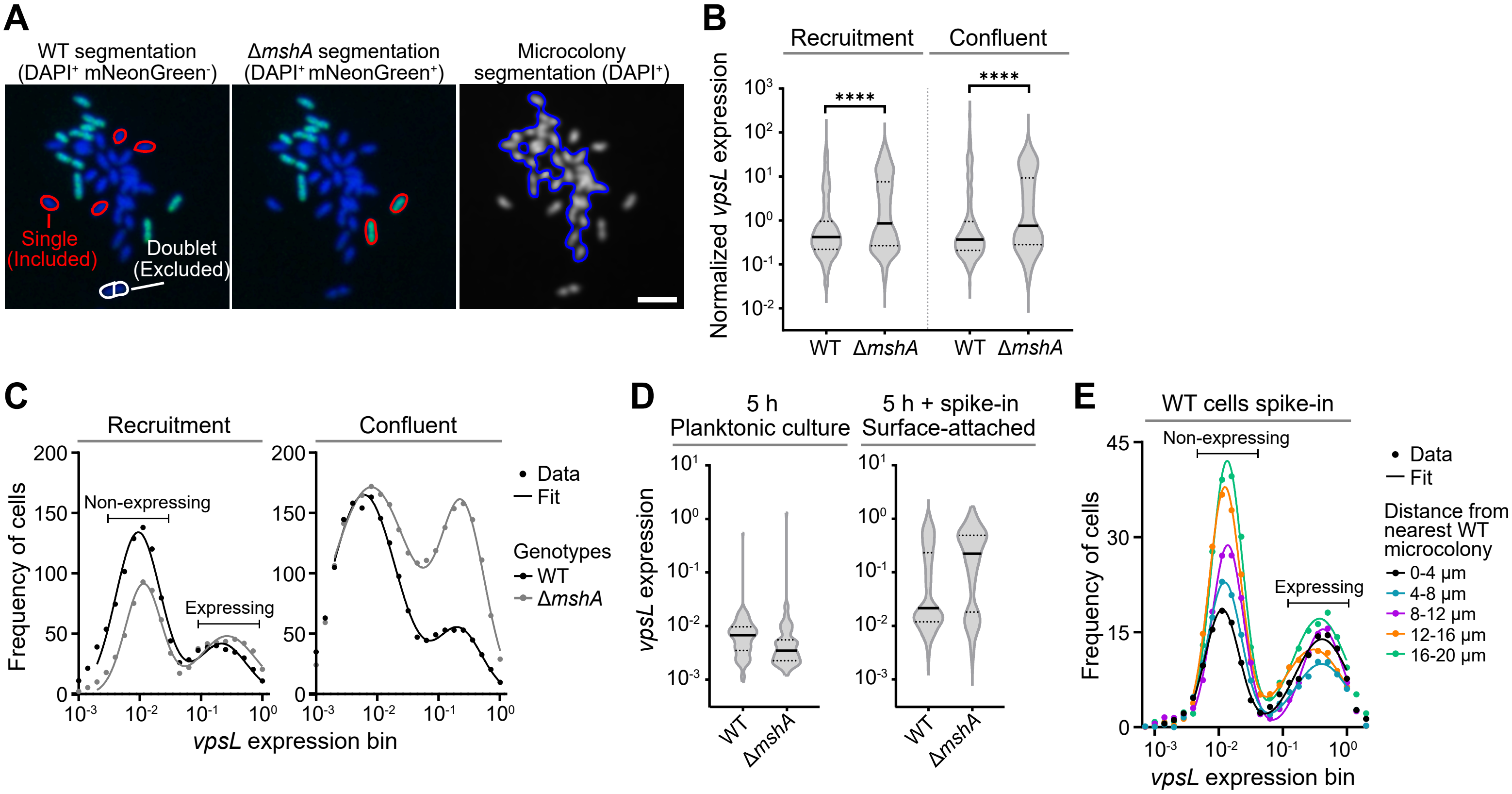


**Figure S5. smFISH quantitation of spatial *vpsL* expression patterns among single recruited cells to regions near existing *V. cholerae* pellicle microcolonies.**

(A) Example image segmentation used to separate genotypes by fluorescence and to separate single cells from microcolonies by morphology. For single-cell segmentation (left and middle images), all cells are labeled in blue (DAPI), and Δ*mshA* cells are additionally labeled in green (mNeonGreen). Red outlines mark cells determined to be single cells through image analysis operations, and the white outline marks a doublet that was excluded from further analysis. For microcolony segmentation (right panel), all cells are labeled in gray (DAPI), and the blue outline indicates a segmented microcolony. Scale bar = 5 μm. (B) *vpsL* expression data shown in main Figure 4B normalized to *gyrA* expression per cell. Solid and dotted black lines are medians and quartiles, respectively. Kolmogorov-Smirnov test ($p_{Recr.}$<0.0001, K-S $D_{Recr.}$=0.25; $p_{Conf.}$<0.0001, K-S $D_{Conf.}$=0.25). Data are from n=6 replicate co-culture wells per stage (Recruitment: WT, n=1119; Δ*mshA*, n=820; Confluent: WT, n=1665; Δ*mshA*, n=2534). (C) Frequency distributions of *vpsL* expression data shown in main Figure 4B. Sum of two Gaussians fits (Table S1). (D) Distributions of per-cell *vpsL* expression among single cells of the designated strains prepared for smFISH following planktonic growth and post-spike-in to cultures containing WT *V. cholerae* pellicle microcolonies at the air-liquid interface. Planktonic data are from n=3 replicates and spike-in data are from n=6 spike-in wells (WT planktonic, n=124; Δ*mshA* planktonic, n=102; WT spike-in, n=3697; Δ*mshA* spike-in, n=541). (E) Frequency distributions of *vpsL* expression data among WT spiked-in cells shown in main Figure 4E with additional distance bins. Sum of two Gaussians fits (Table S1). Data are from n=6 replicate spike-in wells (0-4 μm, n=172; 4-8 μm, n=166; 8-12 μm, n=200; 12-16 μm, n=245; 16-20 μm, n=287).

Table S1. Parameters for Gaussian mixture models fit to smFISH data.

| Sum of two Gaussians | | | | | | | | | |
|---|---|---|---|---|---|---|---|---|---|
| | Figure S5C | | | | Figure S5E | | | | |
| Best-fit values | WT Recruitment | Δ*mshA* Recruitment | WT Confluent | Δ*mshA* Confluent | 0-4 μm | 4-8 μm | 8-12 μm | 12-16 μm | 16-20 μm |
| Amplitude1 | 134 | 90.8 | 165.2 | 171.5 | 18.57 | 22.89 | 28.75 | 37.78 | 42.01 |
| Mean1 | -2.026 | -1.942 | -2.219 | -2.099 | -1.946 | -1.91 | -1.861 | -1.905 | -1.87 |
| SD1 | 0.3666 | 0.2962 | 0.5333 | 0.6586 | 0.2658 | 0.2665 | 0.2447 | 0.2474 | 0.2428 |
| Amplitude2 | 41.77 | 48.12 | 53.17 | 147.2 | 13.9 | 9.995 | 15.45 | 12.29 | 17.15 |
| Mean2 | -0.6607 | -0.5747 | -0.6677 | -0.6094 | -0.3872 | -0.3965 | -0.3736 | -0.5203 | -0.4258 |
| SD2 | 0.3976 | 0.471 | 0.367 | 0.3598 | 0.4132 | 0.4174 | 0.3159 | 0.4693 | 0.4273 |
| Goodness of Fit | | | | | | | | | |
| Degrees of Freedom | 13 | 13 | 13 | 13 | 11 | 11 | 11 | 11 | 11 |
| R squared | 0.9741 | 0.964 | 0.9936 | 0.9892 | 0.9726 | 0.9937 | 0.994 | 0.9959 | 0.9917 |
| Sum of Squares | 699.2 | 337.4 | 302.4 | 257.1 | 10.95 | 3.826 | 6.531 | 6.975 | 17.28 |
| Sy.x | 7.334 | 5.094 | 4.823 | 4.447 | 0.9976 | 0.5898 | 0.7705 | 0.7963 | 1.253 |
| Number of points | | | | | | | | | |
| # of X values | 19 | 38 | 57 | 76 | 17 | 51 | 85 | 119 | 153 |
| # Y values analyzed | 19 | 19 | 19 | 19 | 17 | 17 | 17 | 17 | 17 |

Table S2. Strains used in this study.

| Strain | Identifier | Genotype | Origin |
|---|---|---|---|
| *Vibrio cholerae* C6706 | BB-Vc0045 | *Vibrio cholerae* C6706 | Bassler Lab Collection |
| | BB-Vc0340 | Δ*VC1807*::$P_{tac}$-*mNeonGreen*::*SpecR* | Bassler Lab Collection |
| | BB-Vc0456 | Δ*VC1807*::$P_{tac}$-*mScarlet-I*::*SpecR* | Bassler Lab Collection |
| | BB-Vc0887 | Δ*VC1378*::$P_{tac}$-*ssMBP-AM2-2*::*AmpR* Δ*VC1807*::$P_{luxC}$-*ssMBP-dl5*::*KanR* | Bassler Lab Collection |
| | HKG094 | Δ*mshA* Δ*VC1378*::$P_{tac}$-*ssMBP-AM2-2*::*AmpR* Δ*VC1807*::$P_{luxC}$-*ssMBP-dl5*::*KanR* | This study |
| | HKG063 | Δ*mshA* Δ*VC1807*::$P_{tac}$-*mNeonGreen*::*SpecR* | This study |
| | HKG039 | Δ*mshA* Δ*bap1* Δ*rbmC* Δ*VC1807*::$P_{tac}$-*mScarlet-I*::*SpecR* | This study |
| | HKG040 | Δ*mshA* Δ*vpsL* Δ*VC1807*::$P_{tac}$-*mScarlet-I*::*SpecR* | This study |
| | HKG052 | Δ*mshA* Δ*VC1807*::$P_{luxC}$-*mScarlet-I*::*SpecR* Δ*lacZ*::$P_{tac}$-*mNeonGreen* | This study |
| | HKG080 | $luxO^{D61A}$ Δ*mshA* Δ*VC1807*::$P_{luxC}$-*mScarlet-I*::*SpecR* Δ*lacZ*::$P_{tac}$-*mNeonGreen* | This study |
| | HKG100 | $luxO^{D61E}$ Δ*mshA* Δ*VC1807*::$P_{luxC}$-*mScarlet-I*::*SpecR* Δ*lacZ*::$P_{tac}$-*mNeonGreen* | This study |
| | HKG098 | Δ*mshA* Δ*VC1807*::$P_{BAD}$-*vpsT*::*KanR* Δ*lacZ*::$P_{tac}$-*mNeonGreen* | This study |
| | HKG092 | $luxO^{D61A}$ Δ*mshA* Δ*VC1807*::$P_{BAD}$-*vpsT*::*KanR* Δ*lacZ*::$P_{tac}$-*mNeonGreen* | This study |
| | HKG102 | $luxO^{D61E}$ Δ*mshA* Δ*VC1807*::$P_{BAD}$-*vpsT*::*KanR* Δ*lacZ*::$P_{tac}$-*mNeonGreen* | This study |
| | HKG034 | Δ*bap1* Δ*rbmC* Δ*VC1807*::$P_{tac}$-*mScarlet-I*::*SpecR* | This study |
| | HKG035 | Δ*vpsL* Δ*VC1807*::$P_{tac}$-*mScarlet-I*::*SpecR* | This study |
| | HKG036 | Δ*mshA* Δ*VC1807*::$P_{tac}$-*mScarlet-I*::*SpecR* | This study |
| | HKG070 | Δ*bap1* Δ*rbmC* Δ*VC1807*::$P_{tac}$-*mNeonGreen*::*SpecR* | This study |
| | HKG071 | Δ*vpsL* Δ*VC1807*::$P_{tac}$-*mNeonGreen*::*SpecR* | This study |

**Movie S1. WT *V. cholerae* pellicle formation imaged at single-cell resolution, with an *xy* view of maximum projections in *z* over time (left), enlarged inset (white box, right) and *xz* view (fuchsia box, bottom).**

**Movie S2. Growth of Δ*mshA V. cholerae* at the air-liquid interface following addition of BSA or Bap1$^{\Delta 57aa}$.**

**Movie S3. Attached/swimming single cells at the air-liquid interface for the designated *V. cholerae* mutants following addition of Bap1$^{\Delta 57aa}$.**

**Movie S4. Attached/swimming Δ*mshA V. cholerae* cells (pre-cultured for the designated durations) after spike-in to a culture of WT *V. cholerae* pellicle microcolonies.**

**Movie S5. Movement of WT and Δ*vpsL V. cholerae* cells attached to the air-liquid interface at high spatial and temporal resolution.**

**Dataset S1. Data underlying all figures.**
